# Characterization of transcript enrichment and detection bias in single-nuclei RNA-seq for mapping of distinct human adipocyte lineages

**DOI:** 10.1101/2021.03.24.435852

**Authors:** Anushka Gupta, Farnaz Shamsi, Nicolas Altemos, Gabriel F. Dorlhiac, Aaron M. Cypess, Andrew P. White, Mary Elizabeth Patti, Yu-Hua Tseng, Aaron Streets

## Abstract

Single-cell RNA-sequencing (scRNA-seq) enables molecular characterization of complex biological tissues at high resolution. The requirement of single-cell extraction, however, makes it challenging for profiling tissues such as adipose tissue where collection of intact single adipocytes is complicated by their fragile nature. For such tissues, single-nuclei extraction is often much more efficient and therefore single-nuclei RNA-sequencing (snRNA-seq) presents an alternative to scRNA-seq. However, nuclear transcripts represent only a fraction of the transcriptome in a single cell, with snRNA-seq marked with inherent transcript enrichment and detection biases. Therefore, snRNA-seq may be inadequate for mapping important transcriptional signatures in adipose tissue. In this study, we compare the transcriptomic landscape of single nuclei isolated from preadipocytes and mature adipocytes across human white and brown adipocyte lineages, with whole-cell transcriptome. We demonstrate that snRNA-seq is capable of identifying the broad cell types present in scRNA-seq at all states of adipogenesis. However, we also explore how and why the nuclear transcriptome is biased and limited, and how it can be advantageous. We robustly characterize the enrichment of nuclear-localized transcripts and adipogenic regulatory lncRNAs in snRNA-seq, while also providing a detailed understanding for the preferential detection of long genes upon using this technique. To remove such technical detection biases, we propose a normalization strategy for a more accurate comparison of nuclear and cellular data. Finally, we demonstrate successful integration of scRNA-seq and snRNA-seq datasets with existing bioinformatic tools. Overall, our results illustrate the applicability of snRNA-seq for characterization of cellular diversity in the adipose tissue.

## 1 INTRODUCTION

Adipose tissue is a complex, heterogenous organ responsible for maintaining energy balance in animals, by storing energy during nutritional excess and providing energy during nutritional deprivation. This regulation of whole-body energy homeostasis is primarily maintained by two functionally different types of fat: white adipose tissue (WAT), the primary site of lipid storage, and brown adipose tissue (BAT), which specializes in thermogenic energy expenditure. An imbalance in the metabolic activity or expansion of WAT and BAT is implicated in the pathogenesis of lipodystrophy or obesity and associated comorbidities like cardiovascular diseases and type 2 diabetes (Jo et al. 2009; Carobbio et al. 2011; Levelt et al. 2016). Further complexity arises from the heterogeneity within WAT which also includes a cellular subtype called beige adipocytes with greater oxidative capacity (Pfeifer and Hoffmann 2015). Therefore, understanding the molecular pathways of adipose tissue expansion (adipogenesis) in humans and identifying resident cell types that regulate adipocyte activity is necessary for understanding the tissue’s contribution in the pathology of such metabolic diseases. Consequently, recent studies are beginning to elucidate cellular heterogeneity and developmental pathways in distinct adipose tissue lineages at the single-cell level (Deutsch et al. 2020; Ferrero et al. 2020).

Over the last decade, single-cell RNA-sequencing (scRNA-seq) has proven to be a powerful tool for transcriptomic profiling of complex tissues in an unbiased manner (Birnbaum 2018; Chen et al. 2018; Trapnell 2015). This technological revolution has been facilitated by the development of microfluidic workflows for scRNA-seq that make it possible to analyze hundreds to thousands of single cells in one experiment (Birnbaum 2018), paving the way for the construction of a human cell atlas (Regev et al. 2017). Indeed, multiple recent studies using microfluidic scRNA-seq approaches are investigating the heterogeneity of adipocyte precursors (referred to as preadipocytes in the text) in mice (Rondini and Granneman 2020; Ferrero et al. 2020). However, transcriptomic profiling of single mature adipocytes has been challenging, in part because of the technical barriers associated with isolating intact, single adipocytes. Primary adipocytes can be difficult to work with due to their fragile nature, high buoyancy, and large size (Deutsch et al. 2020). Existing protocols for tissue digestion and single-cell suspension preparation often result in complete or partial adipocyte lysis and therefore are not compatible with scRNA-seq library preparation. Consequently, transcriptomic analysis of adipocytes has relied on bulk RNA-sequencing of clonal cell populations (Shinoda et al. 2015; Xue et al. 2015; Min et al. 2016; Lee et al. 2019; Gao et al. 2017a) or scRNA-sequencing of adipocytes harvested by precise pipetting (Spaethling et al. 2016), making generation of individual clones or isolates the rate limiting step. More recently, microfluidic scRNA-seq was used to identify transcriptomic heterogeneity within murine brown adipocytes (Song et al. 2020), with library preparation limited to adipocytes relatively smaller in size as bigger adipocytes can easily rupture in microchips or during droplet formation. Such size-fractionated application of scRNA-seq, however, results in loss of transcriptional patterns uniquely associated with bigger adipocytes (Blüher et al. 2004). To address the challenge of working with tissues that are difficult to dissociate into single cells, recent studies have turned to single-nucleus RNA-sequencing (snRNA-seq) as an alternative approach for transcriptomic profiling of cellular heterogeneity within primary tissue (Gao et al. 2017b; Lake et al. 2016; Wu et al. 2019; Sathyamurthy et al. 2018; Rajbhandari et al. 2019; Krishnaswami et al. 2016; Lacar et al. 2016; Sun et al. 2020; Habib et al. 2016; Liang et al. 2019; Zeng et al. 2016). These studies rely on nuclear mRNA to serve as a proxy for the single-cell transcriptome, and take advantage of protocols which enable efficient extraction of intact nuclei (Krishnaswami et al. 2016; Habib et al. 2017; Benitez and Shinoda 2020; Nee et al. 2018; Rajbhandari et al. 2019). As a result, recent investigations have already started reporting the existence of multiple adipocyte subtypes in humans using snRNA-seq (Rajbhandari et al. 2019; Sun et al. 2020). However, a single nucleus contains 10-100-fold less mRNA than whole-cells, raising the question whether the composition of mRNA transcripts in the nucleus is sufficient to enable identification of the same cell populations as whole-cells. Previous comparisons of single-cell and single-nucleus approaches suggest that in certain tissues, sampling the nuclear transcriptome is sufficient to characterize cellular composition (Selewa et al. 2020; Wu et al. 2019; Bakken et al. 2018; Habib et al. 2017; Lake et al. 2017). However, collectively these studies also demonstrate that the relationship between nuclear and cytoplasmic mRNA is tissue-specific (Lake et al. 2017; Thrupp et al. 2020). Therefore, there is a need to understand the transcriptomic similarities and differences between single-cell and single-nucleus profiles in the context of the human adipose tissue, for which there is growing need to rely on snRNA-seq.

In this study, we explored the ability of snRNA-seq to recapitulate the transcriptional profiles observed by scRNA-seq in the human adipose tissue white and brown lineages. We focused our study on a well-controlled *in vitro* system of human white and brown adipogenesis (Xue et al. 2015; Kriszt et al. 2017). In this *in vitro* model, paired white and brown primary preadipocytes were isolated from a defined anatomical location (the neck depot) of a single individual. This system allowed us to measure cell-to-cell transcriptomic variations within and between lineages, while controlling for inter-individual variabilities that are typically associated with transcriptomic profiling of primary human adipose tissue, such as body mass index, genotype, and gender. Preadipocytes from both lineages were isolated while preserving their intrinsic cellular heterogeneity and were then immortalized to allow for long-term *in vitro* cell-culture. Previously reported data demonstrated that the preadipocyte populations could be differentiated into mature adipocytes with gene expression profiles that correspond to the adipogenic and thermogenic function of primary tissue from human neck BAT and WAT (Xue et al. 2015). Moreover, the *in vitro* cell-culture system allows for isolation of intact nuclei as well as intact single cells across well-defined stages of adipogenesis including mature, lipid-laden white and brown adipocytes. Using this system, we first mapped the cellular heterogeneity at the preadipocyte stage. Both white and brown preadipocytes were processed using a commercial high-throughput single-cell sequencing platform (10x Genomics). We then extracted nuclei from these populations and performed snRNA-seq using the same isolation and sequencing protocol. We sequenced snRNA-seq libraries to saturation and compared their transcriptomic profiles with those obtained from scRNA-seq across different cellular subtypes. We next developed a single-adipocyte whole-cell isolation protocol and mapped cellular heterogeneity in mature white adipocytes using the molecular single-cell RNA barcoding and sequencing (mcSCRB-seq) protocol (Bagnoli et al. 2018). The transcriptomic profiles obtained were compared with molecular profiles of single nuclei isolated from the same population of adipocytes. Our analyses characterized the accuracy with which snRNA-seq can identify cell types present at the precursor and mature stages of adipogenesis. We identified both technical and biological artifacts that can introduce gene detection biases in snRNA-seq, and we systematically evaluate the limitations of these biases in the context of human adipogenesis. Finally, we propose a normalization strategy for the removal of systematic technical biases between scRNA-seq and snRNA-seq and demonstrate recovery of shared biology by integrating the two datasets using scVI, a variational autoencoder based framework for analysis of scRNA-seq data (Lopez et al. 2018).

## 2 RESULTS

### 2.1 scRNA-seq reveals transcriptional landscape of white and brown preadipocytes

Unsupervised clustering of white and brown preadipocyte scRNA-seq library grouped the cells into three clusters, referred to as populations 0, 1 and 2 (Fig. 1A). White preadipocytes organized into a single homogeneous cell population, cluster 0, whereas brown preadipocytes revealed two cell populations, cluster 1 and cluster 2 (Fig. 1A). All populations were devoid of endothelial (*CD31*) and hematopoietic marker genes (*CD45*, Fig. S2B and S2C) and reflected a preadipocyte state on the basis of their high expression for common mesenchymal stem cell markers *CD29*, *CD90*, *CD44*, and *ENG* (Fig. S2D to S2G). All populations also had positive expression for adipogenesis regulators *CEBPB*, *PPARG*, and *ZEB1*, further verifying an adipogenic fate for these cells (Fig. S2H to S2J).

**Figure 1.**
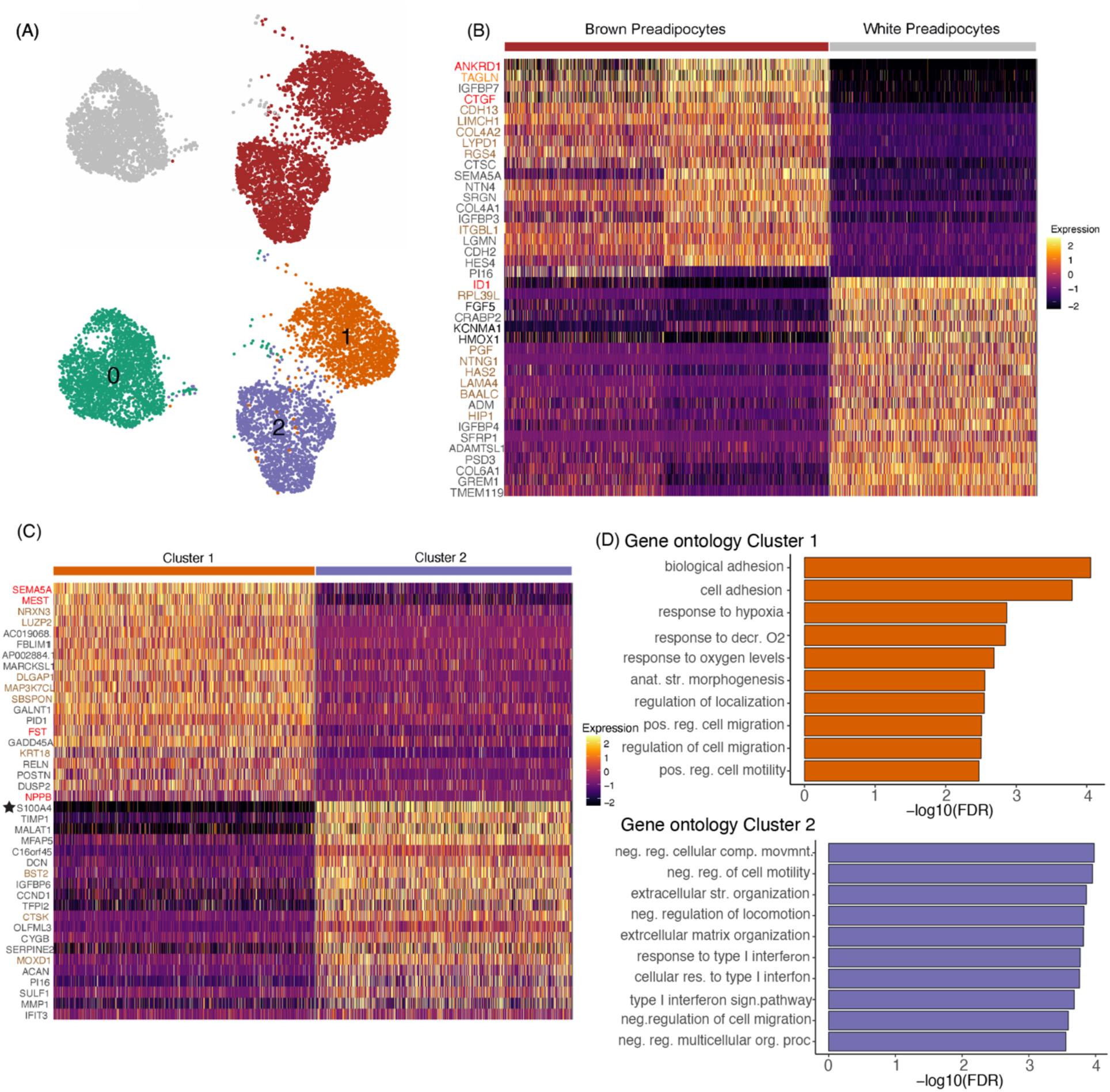
scRNA-seq reveals transcriptional and compositional landscape of white and brown preadipocytes. **(A)** UMAP visualization of white and brown preadipocytes annotated either manually to reflect the sample of origin (top panel) or based on unsupervised clustering (bottom panel) **(B)** Heat map of top 20 differentially expressed genes between white and brown preadipocytes based on log fold-change values. Highlighted in red are regulators of cell proliferation, in orange is smooth-muscle lineage gene, and in beige are genes classified as white- or brown-specific markers (see Note S4). **(C)** Heat map of top 20 differentially expressed genes between brown cluster 1 and cluster 2 based on log fold-change values. Highlighted in red are regulators of adipose development, and in beige are genes classified as cluster-1- or cluster-2-specific markers (see Note S4). **(D)** Top 10 gene ontology biological processes terms enriched in brown preadipocyte cluster 1 (top panel) and cluster 2 (bottom panel). decr. = decreased; anat. str. = anatomical structure; pos. reg. = positive regulation; neg. reg = negative regulation; str. = structure

Differential gene expression (DGE) analysis confirmed that white preadipocytes showed enrichment of genes that are reported to be involved in establishing white preadipocytes’ identity (Table S1B) such as *TCF21* (de Jong et al. 2015), *PAX3* (Sanchez-Gurmaches et al. 2016), and *PDGFRA* (Berry and Rodeheffer 2013). The most upregulated gene in white preadipocytes was *ID1* (Fig. 1B highlighted in red), which is known to maintain progenitor state in preadipocytes by positively regulating the progression of cell cycle for sustained growth and proliferation (Patil et al. 2014; Satyanarayana et al. 2012). Consequently, enriched expression of *ID1* in white preadipocytes suggested ongoing signaling for maintenance of cellular proliferation. In brown preadipocytes, the top upregulated genes included *ANKRD1* and *CTGF* (Fig. 1B highlighted in red), which are well-characterized YAP target genes (Yu et al. 2012). YAP/TAZ are mechanosensitive transcriptional co-activators that regulate proliferation and differentiation at precursor state (Dupont et al. 2011; Hansen et al. 2015; Zhang et al. 2018), while also maintaining thermogenic activity at mature adipocyte state in brown lineage (Tharp et al. 2018). Therefore, our results suggest that brown preadipocytes may have ongoing YAP/TAZ activity for maintenance of brown-lineage progenitor state. DGE analysis also revealed upregulation of smooth-muscle lineage marker genes in brown preadipocytes, such as *TAGLN* (Fig. 1B highlighted in orange), *ACTA2*, *MYL9*, and *CNN1* (Table S1B). These findings are consistent with a recent study that demonstrated abundant expression of smooth muscle lineage–selective genes in clonal human brown preadipocytes (Shinoda et al. 2015), suggesting that brown preadipocytes derived from human neck depot may share this lineage.

Interestingly, we identified two distinct cell populations within brown preadipocytes (cluster 1 and cluster 2, Fig. 1A). Gene ontology (GO) analysis identified cellular adhesion, and regulation of cellular motility as the most enriched terms in cluster 1 (Fig. 1D), suggesting the prevalence of stem-cell-like migratory behavior in these cells. Transforming growth factor superfamily genes (*BMP4* and *TGFB2*) were also enriched in cluster 1 (Table S1C), which play an important role in regulating adipocyte commitment in mesenchymal stem cells (Modica and Wolfrum 2017; Li and Wu 2020). Investigating differential activity of transcription factors (TFs) in cluster 1, transcription factor enrichment analysis (TFEA) identified *FOX* (*FOXC2* and *FOXL1*) and *FOSL1* transcription factors (TFs) with high activity (Table S1D). *FOXC2* participates in the early regulation of preadipocyte differentiation (Gerin et al. 2009; Lidell et al. 2013) while *FOSL1* proteins have been implicated as regulators of cell differentiation, and transformation (Luther et al. 2014, 2011). Therefore, our results indicate that cluster 1 cells may exhibit migratory behavior with ongoing signaling similar to adipogenic fate commitment in mesenchymal stem cells, a behavior we refer to here as stem-cell-like. Enrichment of multiple regulators of adipose tissue development was also detected in cluster 1, such as *SEMA5A* (Giordano et al. 2001), *NPPB* (Villarroya and Vidal-Puig 2013), *MEST* (Karbiener et al. 2015), and *FST* (Braga et al. 2014, Fig. 1C highlighted in red), further suggesting the existence of adipogenic commitment activity in this cell population.

Cluster 2 cells were marked by the expression of *S100A4* gene, also known as the fibroblast specific protein 1 (*FSP1*, Fig. 1C star-marked, Fig. S3D and S3F), which is considered a reliable marker of fibroblasts. GO analysis showed enrichment of immune response, extracellular structure and matrix organization, and negative regulation of cell migration terms in this cell population (Fig. 1D). Multiple genes encoding for extracellular matrix (ECM) components such as *MFAP5*, *ECM1*, *COL6A2*, and *ACAN* were also enriched in cluster 2 (Table S1C). Recent investigations have reported the presence of Fsp1+ fibroblasts in the adipogenic niche, with potential role in maintaining adipose homeostasis (Zhang et al. 2018; Hou et al. 2018; Vijay et al. 2020). The markers identified for fibroblasts in these investigations *FBN1, IGFBP6, MFAP5, FSP1*, and *PI16* were some of the most enriched markers of cluster 2 cells (Fig. S3). Taken together, these results indicate that cluster 2 cells are fibroblast-like, with negative regulation of cellular migration and an ongoing activity for ECM organization.

### 2.2 snRNA-seq identifies the same preadipocyte populations as scRNA-seq and detects biologically relevant differential expression

To evaluate the efficacy of snRNA-seq for recovering transcriptional heterogeneity, we sequenced the nuclear transcriptome of single preadipocytes from the white and brown lineages. Unsupervised clustering of the two lineages grouped nuclei into four clusters, referred to as populations 0, 1, 2 and 3 (Fig. S4A and S4B). Cluster 3 nuclei, however, had enriched expression for stress response genes and mitochondrial genes, along with high background RNA contamination (Fig. S4C), and hence were removed from downstream analyses. In the remaining clusters, brown nuclei were primarily grouped into clusters 1 and 2 whereas white nuclei grouped into a single cluster 0 (Fig. 2A). Similarity between clusters identified in snRNA-seq and scRNA-seq was assessed using the concept of transcriptional signatures (Gaublomme et al. 2019; DeTomaso and Yosef 2016), defined as genes differentially expressed in either white vs brown preadipocytes, or cluster 1 vs cluster 2, in the scRNA-seq dataset (Table S1B and S1C). As expected, the transcriptional signature scores, calculated using Vision (DeTomaso et al. 2019), were enriched in the corresponding preadipocyte-type/clusters in the snRNA-seq dataset (Fig. 2B), thereby demonstrating a high concordance between transcriptional features uncovered by the two techniques.

**Figure 2.**
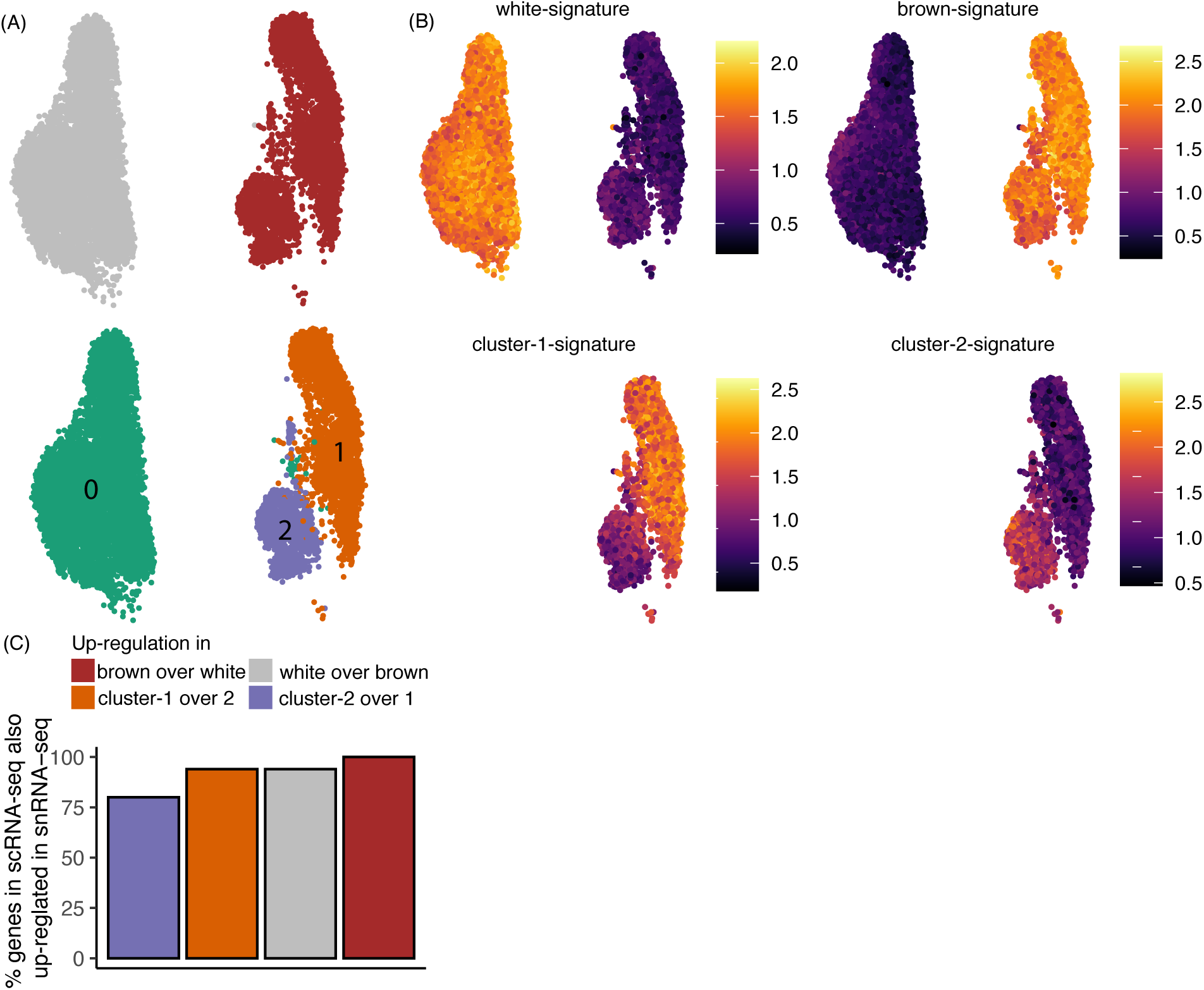
snRNA-sequencing identifies the same preadipocyte populations as scRNA-seq and detects biologically relevant differential expression. **(A)** UMAP visualization of white and brown preadipocytes annotated either manually to reflect the sample of origin (top panel) or based on unsupervised clustering (bottom panel) **(B)** Heatmap of transcriptional signature scores for white preadipocyte (top left panel), brown preadipocyte (top right panel), brown preadipocyte cluster 1 (bottom left panel), and brown preadipocyte cluster 2 (bottom right panel) as plotted on the UMAP visualization of snRNA-seq data **(C)** Bar plot of percent top-50 genes differentially enriched (DE) in scRNA-seq dataset that are also DE in snRNA-seq dataset. Top-50 genes were evaluated based on log fold-change values using scRNA-seq dataset.

As was observed with scRNA-seq, white nuclei were enriched for genes *TCF21, PAX3* and *PDGFRA* (Table S2A), and brown nuclei were enriched for YAP/TAZ target genes *ANKRD1* and *CTGF* (Table S2A), and smooth muscle lineage marker genes *TAGLN, MYL9, CNN1*, and *MYH11* (Table S2A). Gene *ID1*, however, was not differentially enriched in white nuclei. The inability to detect differential enrichment of *ID1* in nuclei could be caused by limited detection in snRNA-seq. In scRNA-seq dataset, we had classified certain DE genes as markers for white and brown preadipocytes based on their highly enriched and specific expression (Note S4). All such white- and brown-preadipocyte specific marker genes were also enriched in white and brown nuclei respectively (Table S2A). Of the 50 genes with maximum enrichment (ordered by logFC) in white and brown preadipocytes in scRNA-seq dataset, over 94% were also differentially expressed in white and brown nuclei respectively (Fig. 2C). This analysis demonstrates that snRNA-seq has sufficient sensitivity to recover same molecular differences as scRNA-seq between white and brown preadipocytes.

GO analysis identified enrichment of cellular adhesion, and regulation of cellular localization terms in brown cluster 1 nuclei, corresponding with the findings in scRNA-seq dataset (Fig. S4D). Transforming growth factor superfamily genes *BMP4* and *TGFB2* were also enriched in cluster 1, along with regulators of adipose tissue development *SEMA5A*, *MEST*, and *FST* (Table S2B). All 6 cluster-1-specific marker genes (Note S4) identified were also enriched in cluster 1 nuclei (Table S2B). Of the 50 genes with maximum enrichment (ordered by logFC) in cluster 1 cells in scRNA-seq dataset, 94% were also differentially expressed in the nuclear dataset (Fig. 2C). In cluster 2 brown nuclei, enrichment of *FSP1* was observed (Table S2B), as well as regulation of extracellular matrix organization terms based on GO analysis (Fig. S4E). Genes encoding for extracellular matrix components *COL6A2, MFAP5, ACAN*, and *ECM1* were all upregulated in cluster 2 (Table S2B). Of the 50 genes with maximum enrichment (logFC) in cluster 2 brown preadipocytes (scRNA-seq dataset), 80% were also differentially expressed in the nuclear dataset (Fig. 2C). All cluster-2-specific marker genes (Note S4) identified were also enriched in cluster 2 nuclei (Table S2B). Overall, our snRNA-seq analyses indicated the emergence of stem-cell-like behavior in cluster 1 and fibroblast-like behavior in cluster 2, in agreement with the whole-cell dataset.

### 2.3 Gene length-associated detection bias in single-nuclei RNA-sequencing

Typical scRNA-seq data analysis pipelines often filter intronic reads for downstream count matrix generation. More recently, however, evidence has suggested that intronic reads originate from nascent transcripts (Ameur et al. 2011; Gray et al. 2014; Hendriks et al. 2014), and hence are informative about expression levels in single-cell data. Furthermore, the additional read counts improve gene detection sensitivity and can improve cell-cluster resolution (Bakken et al. 2018; Wu et al. 2019). Multiple recent studies have suggested internal hybridization of polyT RT-primer to intronic polyA stretches in nascent transcripts as the primary mechanism for the capture and detection of intronic reads (La Manno et al. 2018; Patrick et al. 2020; Shulman and Elkon 2019). Consequently, intronic reads are more readily detected in genes with more intronic polyA stretches, which are more likely to be longer in length (Fig. S1A). This bias is increased in nuclear libraries where up to 40% of all the reads map to intronic regions as compared to only 9% in scRNA-seq (Fig. 3A). Consequently, recent studies have reported enrichment of longer genes (Bakken et al. 2018; Lake et al. 2017) and poor detection of shorter genes (Thrupp et al. 2020) in nuclei.

**Figure 3.**
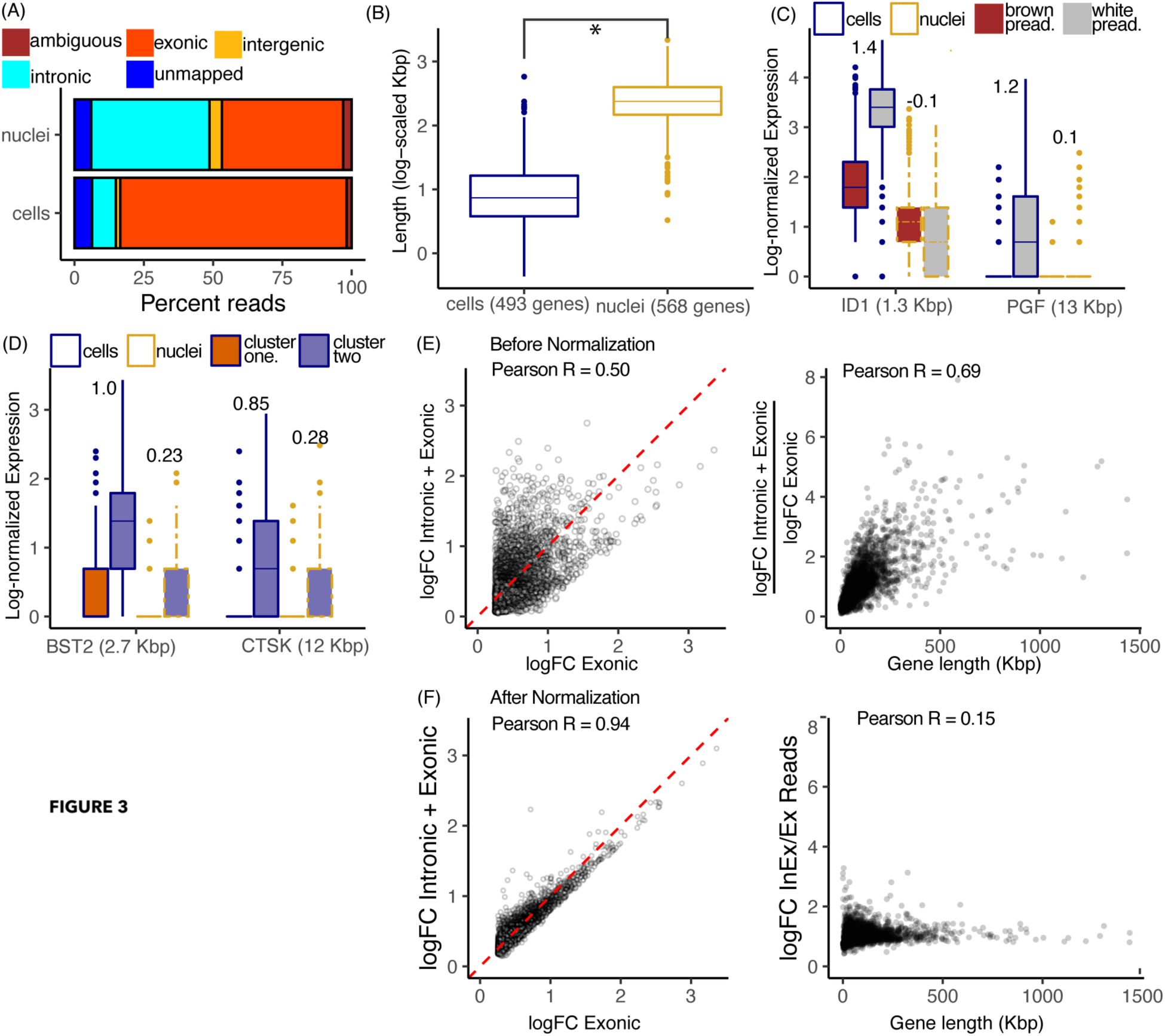
Gene length associated detection bias in the nuclear transcriptome. **(A)** Distribution of percent exonic, intronic, intergenic, ambiguous and unmapped reads in scRNA-seq and snRNA-seq preadipocyte datasets **(B)** Distribution of gene length for genes enriched in cells (in blue) and nuclei (in yellow) with log fold-change > 1 and FDR < 0.05 including both intronic and exonic reads **(C)** Box plot of log library-size-normalized expression values for genes *ID1* and *PGF* in scRNA-seq and snRNA-seq datasets. Black text indicates logFC value for white vs. brown DE test with FDR < 0.05. **(D)** Box plot of log library-size-normalized expression values for genes *BST2* and *CTSK* in scRNA-seq and snRNA-seq datasets. Black text indicates logFC value for cluster 2 vs. cluster 1 DE test with FDR < 0.05.**(E)** Left panel: Log-fold-change for nuclear-enriched genes when using only exonic reads, or both intronic and exonic reads before normalization. Each dot represents a gene enriched in nuclei using exonic-only reads with logFC > 0.25 and FDR < 0.05. Red dotted line indicates y=x axis. Right panel: Ratio of y-axis-value over x-axis-value for genes in left panel, plotted as a function of their length. **(F)** Left panel: Log-fold-change for nuclear-enriched genes when using only exonic reads, or both intronic and exonic reads after normalization. Each dot is the same as in panel **(E)**. Red dotted line indicates y=x axis. Right panel: Ratio of y-axis-value over x-axis-value for genes in left panel, plotted as a function of their length.

To examine the enrichment of long genes in nuclei, we first performed DGE analysis between cells and nuclei in white preadipocytes. Using both intronic and exonic reads, our analysis identified 493 genes enriched in cells and 568 genes enriched in nuclei (logFC >1and FDR <0.05). Notably, nuclear-enriched genes were significantly longer than genes enriched in whole-cells (two-group Mann–Whitney U-test, p-value < 0.01, Fig. 3B). Moreover, single-nuclei measurements revealed poor detection of shorter genes such as white preadipocyte-specific marker genes *ID1* & *PGF* (Fig. 3C) and cluster 2-specific marker genes *BST2* & *CTSK* (Fig. 3D).

We also performed DGE analysis between white cells and white nuclei using only exonic reads (logFC >0.25 and FDR <0.05). Notably, the logFC differential enrichment for nuclear-enriched genes was poorly correlated with counting exons or exons and introns (Fig. 3E, Pearson R = 0.50, p-value < 0.01). logFC values for some of the longest genes were artificially inflated, possibly because of their preferential detection upon inclusion of intronic reads (Fig. 3E Right panel). Conversely, logFC values for some of the shortest genes were artificially deflated because of their poor detection (Fig. 3E Right panel). Consequently, the ratio of the logFC values with counting exons or exons and introns, was strongly correlated with gene length (Fig. 3E, Pearson R = 0.69, p-value < 0.01). Overall, our results demonstrate technical artifacts induced by gene-length associated detection bias in snRNA-seq, upon inclusion of intronic reads. We therefore developed a normalization strategy to address this technical, length-associated detection bias (Note S3). After normalization, the logFC differential enrichment of nuclear-enriched genes was highly correlated with counting exons or exons and introns (Fig. 3F, Pearson R = 0.94, p-value < 0.01). Moreover, the ratio of the logFC values with counting exons or exons and introns, after normalization, was poorly correlated with gene-length (Fig. 3F, Pearson R = 0.15, p-value < 0.01). Nuclear and cellular transcriptomes were also better correlated after removal of technical biases using our normalization strategy (Fig. S5A).

DGE analysis, between white cells and white nuclei, with normalized read counts identified 382 enriched genes in cells and 249 enriched genes in nuclei (logFC >1and FDR <0.05), with nuclear-enriched genes still significantly longer than whole-cells (two-group Mann–Whitney U-test, p-value < 0.01, Fig. S5B). However, the genes enriched in nuclei were on average 14-fold longer than genes enriched in cells (as compared to 32-fold difference before normalization), which is comparable to the difference observed when using only exonic reads (11-fold difference, Fig. S5C), suggesting that after accounting for technical bias, there also exists biological enrichment of longer genes in nuclei. Overall, our observations demonstrate that length-normalization removes artificial detection biases thereby improving UMI count estimation accuracy, while also preserving improved gene detection sensitivity afforded by inclusion of intronic reads.

To further understand differential transcript enrichment between whole-cell and nuclear transcriptomes, we next focused on genes enriched in whole-cells after normalization. GO analysis identified protein translation associated terms as most enriched in whole-cells (Fig. S5D). Genes contributing to the enrichment of translational terms primarily included mRNAs encoding for ribosomal proteins. This enrichment of ribosomal-protein mRNAs in whole-cells is consistent with their very low cytoplasmic decay rates and selective nuclear export machinery (Wickramasinghe et al. 2014; Chen and van Steensel 2017). Yet, poor detection of ribosomal proteins in the nuclear transcriptome did not affect the ability to resolve cellular populations in snRNA-seq data, as evident by the score of transcriptional signature consisting of top 100 genes enriched in cells based on logFC values (∼ 53/100 ribosomal protein genes; Fig. S5E).

### 2.4 Nuclear transcriptome is enriched for long non-coding RNAs that regulate adipogenesis and drive cell-type differences

Long non-coding RNAs (lncRNAs) function in regulating diverse biological processes, including regulation of transcription, proliferation, pluripotency, and cellular differentiation (Quinodoz and Guttman 2014; Sherstyuk et al. 2018; Samata and Akhtar 2018; Delá et al. 2019). Because of their regulatory function, lncRNAs predominantly remain localized in the nucleus (Cabili et al. 2015; Wen et al. 2018). snRNA-seq intrinsically enriches for nuclear localized transcripts, and previous studies have reported enrichment of lncRNAs in snRNA-seq libraries over scRNA-seq (Grindberg et al. 2013; Zeng et al. 2016). We hypothesized that nuclear enrichment of lncRNAs could be advantageous for characterizing adipose tissue because multiple lncRNAs have also been implicated in regulating adipogenesis (Sun et al. 2013; Ding et al. 2018; Wei et al. 2016; Sun and Lin 2019; Zhou et al. 2020). We tested this hypothesis in our *in vitro* system by profiling adipogenic regulatory lncRNAs in our whole-cell and nuclear libraries derived from white preadipocytes, after normalization. We identified significant enrichment of lncRNAs *NEAT1* (Wei et al. 2016), *MEG3* (Li et al. 2017), *MIR31HG* (Huang et al. 2017), and *PVT1* (Zhang et al. 2020) in white nuclei, which are previously reported regulators of adipogenesis (Fig. 4A). All four lncRNAs were also enriched in brown nuclei as compared to brown whole-cells (Fig. S6A to S6D). Generally, snRNA-seq consistently detected a greater number of lncRNAs at all read depths than scRNA-seq (Fig. 4B, p-value < 0.01, two-group Mann–Whitney U-test). Of the 111 differentially expressed lncRNAs between white nuclei and white cells, ∼86% (96/111 genes) were upregulated in nuclei, thereby validating a higher prevalence of this class of genes in the nuclear compartment. 7 out of 15 lncRNAs that were enriched in white cells were snoRNA host genes (SNHGs), that have been shown to have various functions in cytoplasm such as repressing mRNA translation, miRNA sponging, and protein ubiquitination (Zimta et al. 2020). Overall, our results suggest a higher likelihood to deconstruct the functional roles of adipogenic regulatory lncRNAs (and other lncRNAs in general) using snRNA-seq.

**Figure 4.**
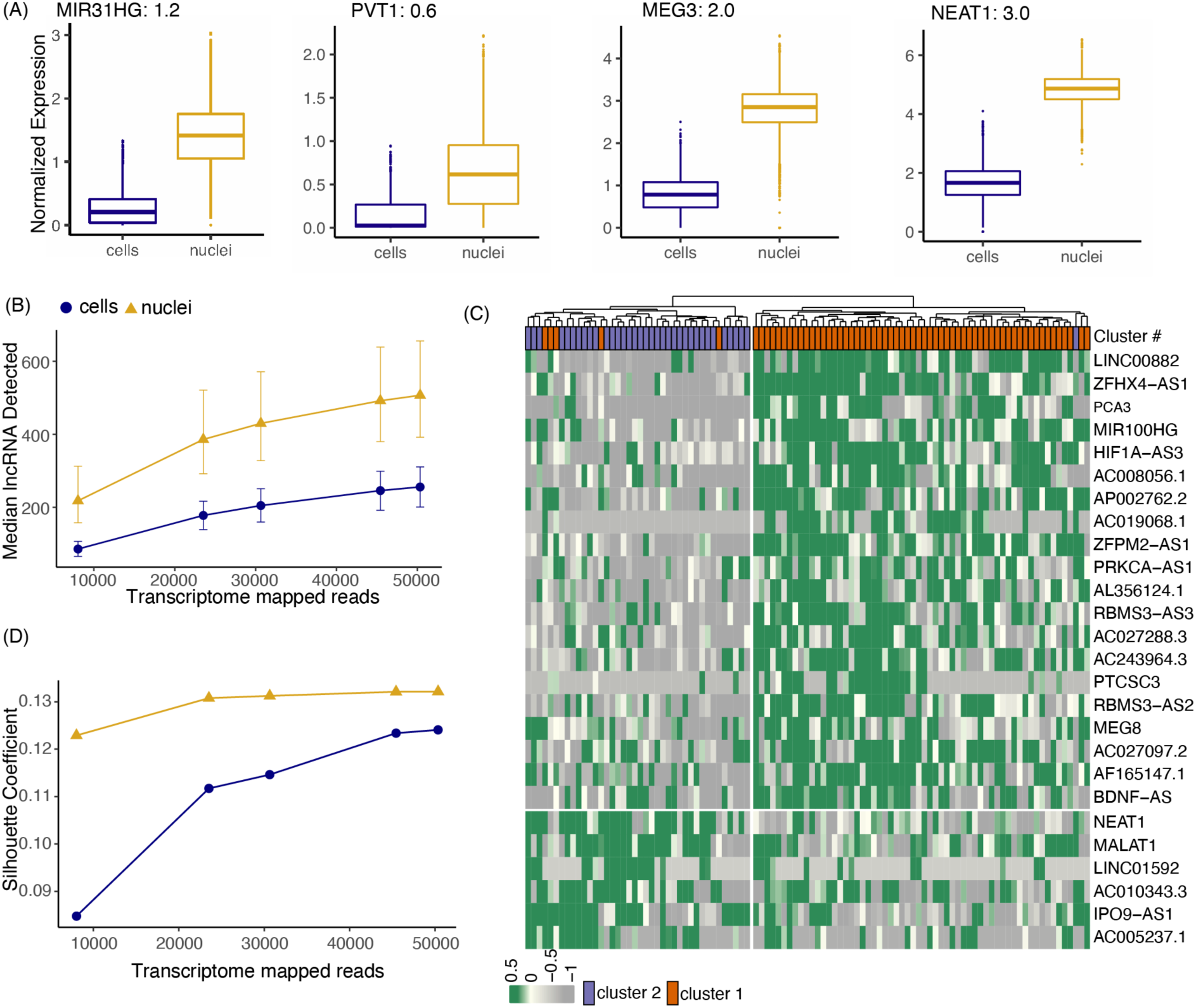
Nuclear transcriptome is enriched for lncRNAs that regulate adipogenesis and drive cell-type differences. **(A)** Boxplots of lncRNAs reported as regulators of adipogenesis. Black text indicates logFC value for white nuclei vs. white cell DE test in preadipocytes with FDR < 0.05 after normalization **(B)** Median lncRNAs detected as a function of read depth across single cells and nuclei (both white and brown lineages). Error bars indicate the interquartile range **(C)** Hierarchical clustering using scaled expression values of top-20 upregulated lncRNAs in brown cluster 1 and cluster 2 in snRNA-seq dataset. 100 random barcodes were chosen for this analysis. Topmost row reflects original cluster assignment for the selected barcodes **(D)** Cluster separation resolution quantification between brown cluster 2 vs cluster 1 in scRNA-seq and snRNA-seq dataset. Only lncRNAs were considered for PCA manifold generation. Both datasets were subsampled to have the same number of cells/nuclei and same number of mean transcriptome mapped reads.

Next, we evaluated the sensitivity of snRNA-seq for detection of lncRNAs driving molecular heterogeneity between brown preadipocyte cluster 1 and 2, two cell-types most closely related to each other. At ∼50,000 reads per cell/nuclei, DGE analysis identified over 40 lncRNAs distinctively regulated between cluster 1 and 2 in the snRNA-seq dataset as compared to only 15 lncRNAs in scRNA-seq dataset. Unsupervised hierarchical clustering in the snRNA-seq dataset based on the expression of top 20 upregulated lncRNAs in cluster 1 and 2 each revealed sorting of nuclei into two distinct groups that predominantly reflected their original cluster assignment (Fig. 4C). Moreover, Silhouette coefficient analysis (a method for evaluating clustering performance) revealed better cluster separation performance for snRNA-seq as compared to scRNA-seq between cluster 1 and 2 for all downsampled read depths (Fig. 4D). Silhouette coefficients were calculated based on Euclidean distance between cells/nuclei in the principal component space generated using only lncRNAs (see Methods). To validate that the observed performance features were not metric dependent, we quantified two more indices, the Calinski-Harabasz Index, and the Davies-Bouldin Index to compute inter-cluster separation and found similar trends (Fig. S6F and S6G). A similar analysis performed by normalizing for the same number of mean unique molecules (UMI) per sample revealed a similar trend for the three separation indices (Fig. S6H to S6J). Together, our results suggest that snRNA-seq is superior for learning heterogeneity governed by lncRNAs as compared to scRNA-seq.

### 2.5 snRNA-seq detects relevant transcriptional regulation during adipogenesis in white preadipocytes

After identifying transcriptomic similarities and differences between scRNA-seq and snRNA-seq in preadipocyte state, we next focused on evaluating molecular correspondence between the two techniques in mature adipocytes. We leveraged our *in vitro* model of white adipogenesis that enabled us to prepare a single-cell suspension of mature adipocytes without the need of implementing harsh tissue dissociation protocols (see Methods). Following single-cell suspension preparation, one of the most common ways to sort single cells is using flow cytometry. Recently, FACS gating strategies have been tailored to isolate mature adipocytes (Hagberg et al. 2018; Majka et al. 2014), although only a small percentage of adipocytes are able to survive the shear stress associated with flow sorting (Majka et al. 2014). Therefore, to enable gentle sorting of single adipocytes for downstream scRNA-seq, we developed a new protocol using the cellenONE® X1 single-cell isolation platform. This automated liquid-handling robot uses gentle piezo-acoustic technology for dispensing cells encapsulated in a picoliter-volume droplet, ensuring minimal cellular perturbation and background RNA contamination. To harvest adipocytes *in vitro*, human white preadipocytes were cultured and differentiated using a chemical adipogenic induction cocktail for 20 days (Shamsi and Tseng 2017). Coherent anti-stokes Raman imaging established successful differentiation of white preadipocytes, with distinctly visible signal from round lipid droplets (Fig. S7A). After creating a single-cell suspension of white adipocytes, 200 cells were spotted using the cellenONEX1 machine, onto 96-well plates preloaded with lysis buffer and barcoded polyT primer. Library preparation was then performed using the mcSCRB-seq chemistry (Bagnoli et al. 2018). Transcriptomic profiles of these cells were then compared with a snRNA-seq library of ∼12,000 nuclei isolated from 20-days differentiated white adipocytes.

Independent unsupervised clustering revealed organization of both cells and nuclei into primarily two clusters, referred to as cluster 0 and 1 (Fig. 5A). snRNA-seq identified an additional cluster 2, which exhibited characteristics of mitotic preadipocytes with ongoing cell cycle progression, suggesting that these cells could be preadipocytes that never underwent growth arrest (Note S2; Fig. S8). Cluster 0 in both datasets was marked by the expression of mesenchymal marker *THY1* (Fig. 5A), suggesting that these cells/nuclei were differentiating preadipocytes. Cluster 1, on the other hand, had high expression of adipogenic gene *ADIPOQ*, indicating that cells/nuclei in this cluster were mature adipocytes (Fig. 5A). DGE analysis further identified enrichment of other adipogenic marker genes (along with *ADIPOQ*) in cluster 1 (Fig. 5B and 5C, highlighted in red), confirming a transition from differentiating preadipocytes to mature adipocytes from cluster 0 to cluster 1 in both datasets. GO analysis identified enrichment of extracellular matrix organization terms in cluster 0 and lipid metabolism in cluster 1, independently in both scRNA-seq and snRNA-seq datasets (Fig. S7B to S7E). Moreover, ∼80% genes (106/133) upregulated in cluster 1 in the scRNA-seq dataset, were also differentially expressed in the snRNA-seq dataset. Notably, the remaining 20% genes (27/133) that were not differentially expressed in the snRNA-seq dataset primarily included genes associated with the mitochondrial respiratory chain process (Fig. S7F), suggesting that adipocytes’ enhanced mitochondrial activity may not be captured in the snRNA-seq dataset. Correspondingly, snRNA-seq dataset lacked manifestation of mitochondrial biological processes such as oxidative phosphorylation, and electron transport chain in cluster 1 upon GO (Fig. S7B vs S7D). This observation was also supported by the fact that these 27 genes had a median length of ∼11 Kbp, the same order of magnitude as length of genes with poor detection in nuclei over whole cells (Fig. 3B). As expected, scores of clusters 0 and 1 transcriptional signatures in the snRNA-seq dataset were observed to be highly conserved and enriched in corresponding cluster types (Fig. 5D and 5E), further validating the conservation of information in the nuclear transcriptome. Overall, our results reveal a comparable molecular landscape in white adipocytes between scRNA-seq and snRNA-seq datasets.

**Figure 5.**
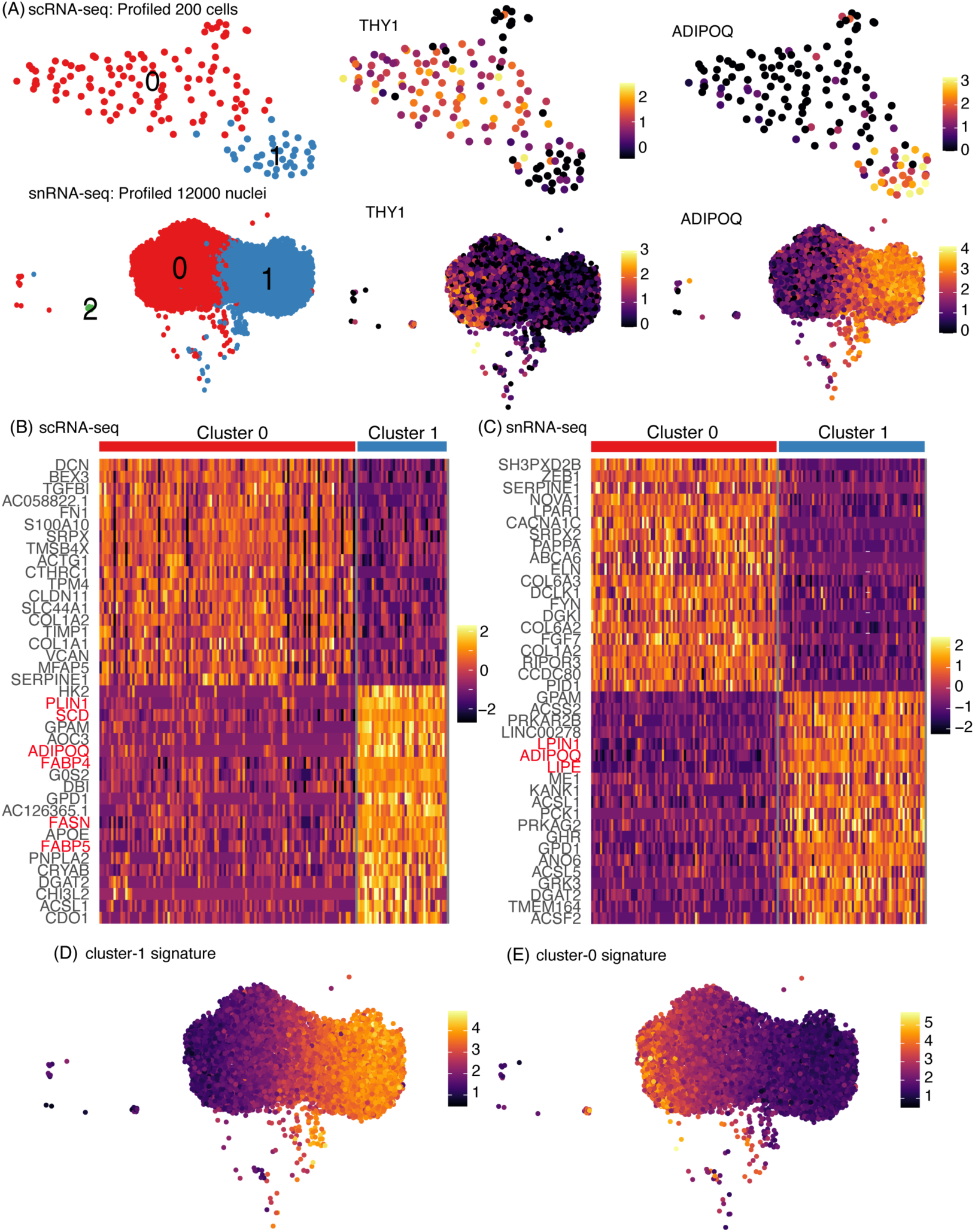
snRNA-seq detects important transcriptional regulation during adipogenesis in white preadipocytes. **(A)** UMAP visualization of scRNA-seq and snRNA-seq white adipocyte datasets (day-20) after unsupervised clustering (leftmost panels). Expression profile for mesenchymal marker *THY1* and mature-adipocyte marker *ADIPOQ* in both scRNA-seq and snRNA-seq datasets (middle and rightmost panels) **(B)** Heat map of z-scored expression of top 20 differentially expressed genes between cluster 0 and cluster 1 in scRNA-seq white adipocyte dataset. Highlighted in red are markers of adipogenesis **(C)** Heat map of z-scored expression of top 20 differentially expressed genes between cluster 0 and cluster 1 in snRNA-seq white adipocyte dataset. A random subset of 150 barcodes was used for this visualization. Highlighted in red are markers of adipogenesis **(D)** Heatmap of transcriptional signature scores for cluster 1 as plotted on the UMAP visualization of snRNA-seq white adipocyte data **(E)** Heatmap of transcriptional signature scores for cluster 0 as plotted on the UMAP visualization of snRNA-seq white adipocyte data

### 2.6 Integration of snRNA-seq and scRNA-seq datasets

A comprehensive cell atlas of the adipose tissue will require joint analyses of datasets generated using both scRNA-seq and snRNA-seq. However, technical biases and differential transcript enrichment in snRNA-seq leads to significant batch effects between snRNA-seq and scRNA-seq experiments, thereby reducing clusterability of cells from these two protocols (Mereu et al. 2020). Multiple bioinformatic tools are now available to remove covariates that lead to technical batch effects and facilitate integration of scRNA-seq datasets generated across different days, laboratories, individuals, or technologies (Zappia et al. 2018). We used single-cell variational inference (scVI), a deep generative modeling-based tool (Lopez et al. 2018), to explore the possibility of integrating snRNA-seq and scRNA-seq datasets for joint analysis. Four datasets of white preadipocytes were integrated in total: day-0 scRNA-seq & snRNA-seq (cluster 0 in Fig. 1A and Fig. 2A), and day-20 scRNA-seq & snRNA-seq (top and bottom left panels in Fig. 5A).

Without batch correction, all four datasets arranged into distinct individual clusters, with no shared population identified at the same time point across different techniques, or same technique but across different time-points (Fig. 6A). A dendrogram, based on the Euclidean distance in dimensionally reduce space, grouped clusters first by sequencing chemistry (mcSCRB-seq vs 10x), followed by technique type (snRNA-seq vs scRNA-seq), and finally by time point (day-0 vs day-20, Fig. 6A). After integration, matching adipocyte populations from day-20 and preadipocyte populations from both day-0 and day-20 in nuclear and whole-cell datasets were primarily nearest neighbors in a dendrogram based on the Euclidean distance in dimensionally reduced space (Fig. 6B). UMAP visualization further revealed proximal placements of similar cell populations (Fig. 6B). Of note, we observed that preadipocytes from both day-0-snRNA-seq and day-0-scRNA-seq datasets localized into two distinct groups, which was driven by differences in proliferation state with one cluster composed of mitotic cells and another composed of growth arrested cells (Note S2; Fig. S8). Overall, our results demonstrate scVI’s integration abilities by identifying functionally similar preadipocyte and adipocyte populations shared across different techniques.

**Figure 6.**
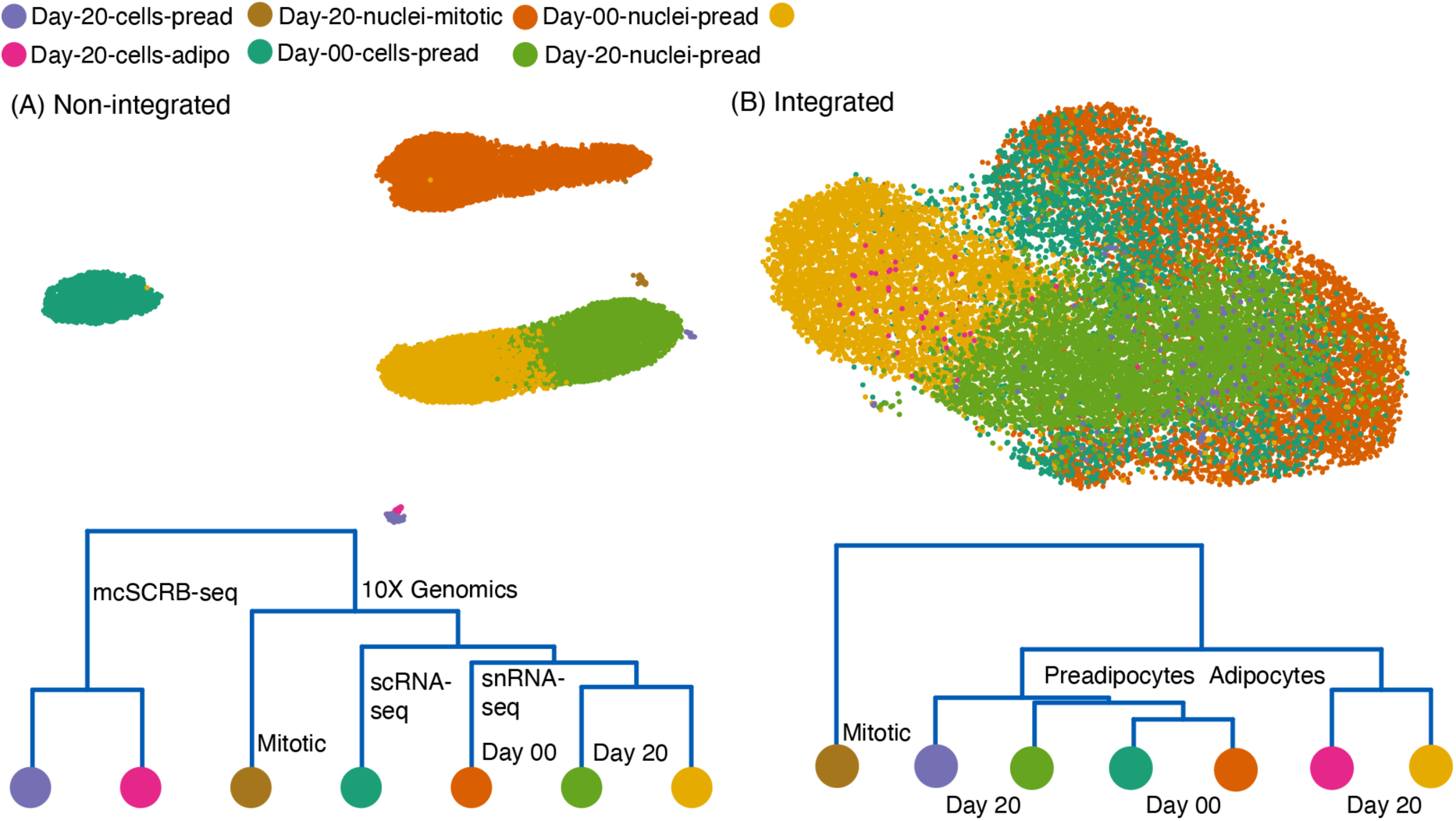
Integration of snRNA-seq and scRNA-seq datasets. **(A)** UMAP visualization of non-integrated scRNA-seq and snRNA-seq datasets for both white preadipocyte (day-0) and mature adipocyte (day-20), for a total of 4 batches (top panel). Cluster dendrogram for non-integrated datasets based on the eigenvalue-weighted Euclidean distance matrix constructed in latent-dimension space inferred using scVI (bottom panel) **(B)** UMAP visualization and cluster dendrogram of scRNA-seq and snRNA-seq datasets as in panel A after integration using scVI-tools. See also Note S2 and Fig. S8.

## 3 Discussion

In this investigation, we evaluated the ability of snRNA-seq to recapitulate the molecular and compositional landscape of distinct lineages in human adipose tissue. We avoided confounding variability associated with inter-depot and inter-subject transcriptional variation by performing a direct comparison of snRNA-seq and scRNA-seq on a pair of immortalized white and brown human preadipocytes isolated from the neck region of the same individual. We found that snRNA-seq was able to recover the same cell-types as scRNA-seq at both preadipocyte and mature adipocyte states. Furthermore, we provided evidence for recovering similar expression profiles of biologically relevant genes, and attributing similar functional annotations to cell-types by nuclear transcriptome profiling as compared to whole-cells.

At the preadipocyte stage, brown preadipocytes were a heterogeneous mix of two distinct cell populations, cluster 1 and cluster 2. However, cell-type enrichment followed by differentiation and metabolic assays will need to be further performed to identify their individual functions in maintaining adipose tissue homeostasis. To date, different scRNA-seq studies of mouse stromal vascular fraction have identified multiple subpopulations of adipose progenitor cells (APCs) expressing distinct markers (Burl et al. 2018; Hepler et al. 2018; Merrick et al. 2019; Schwalie et al. 2018). Integrated analysis of these datasets primarily identified two common populations of APCs in mice referred to as Asc1 and Asc2 (Ferrero et al. 2020; Rondini and Granneman 2020). Similar to cluster 2 cells, Asc2 exhibited pro-inflammatory and pro-fibrotic phenotype and positive expression of genes *PI16* and *MFAP5*. Functional investigations into the two cell types revealed Asc2 cells inhibiting the differentiation of Asc1 cells *in vitro* (Rondini and Granneman 2020). Therefore, it is plausible that cluster 1 and cluster 2 cells identified in our study may be functioning in a manner similar to Asc1 and Asc2 to maintain adipocyte turnover.

snRNA-seq is the preferred technique to study samples whose compositional landscape may be biased by the differential efficiency of cell-type recovery when using scRNA-seq. Adipose tissue is one such sample where isolation of intact, single adipocytes is complicated by their fragile nature. Here, we developed a new single-adipocyte isolation protocol using piezo-acoustic-based gentle dispensing technology for improved recovery with downstream scRNA-seq. However, this adipocyte capture efficiency was still limited as compared to snRNA-seq where ∼ 48% barcodes were identified as adipocytes as compared to only ∼26% in scRNA-seq. Conversely, at the preadipocyte stat, where cell-type recovery is efficient, scRNA-seq recovered equal proportions of the two brown preadipocyte clusters. However, analysis of snRNA-seq data revealed ∼1.5-fold enrichment of cluster 1 over cluster 2, suggesting a bias in compositional sampling in snRNA-seq. Therefore, such cell-level sampling biases must be considered when evaluating the composition of complex tissues with snRNA-seq.

Understanding the advantages and drawbacks of using snRNA-seq, a nuclear transcriptome is inherently enriched for nascent transcripts, thereby predominantly reflecting changes in gene expression as a result of differences in transcription rates alone (Gaidatzis et al. 2015). In contrast, a cellular transcriptome is fundamentally enriched for mature transcripts, thereby capturing gene expression changes driven by both transcriptional and post-transcriptional regulatory processes such as mRNA processing and degradation. Higher relative proportion of nascent to mature transcripts in the nucleus also results in a large fraction of intronic reads in snRNA-seq, which when considered for count matrix generation, gives rise to detection bias against short genes with few intronic polyA stretches. Consequently, for a biological system compatible with both techniques, scRNA-seq may be better for identifying cellular subpopulations. scRNA-seq will also be better for assessing gene expression changes as a result of post-transcriptional regulation. However, nuclear transcriptome is preferentially enriched for lncRNAs, indicating that functional investigations of these genes will be enhanced by sequencing nuclei. Moreover, some studies of specific nuclear functions may be enhanced by directly accessing nuclei for example, changes in gene expression profile as a result of targeted transcriptional activation mediated by epigenetic modifications. Therefore, it is important to evaluate each approach depending on the task at hand. However, for tissues such as the adipose tissue, snRNA-seq may be the only option. In our investigation, lncRNAs regulating adipogenesis were enriched in the nuclear transcriptome. lncRNAs driving differences between cluster 1 and 2 in brown preadipocytes were also better detected in the snRNA-seq dataset. However, we also identified poor detection of shorter genes in nuclei, some of which were key to driving heterogeneity between distinct cell-types.

Including intronic reads for UMI quantification presents researchers with both advantages and drawbacks. polyA stretches are found randomly dispersed along the length of the genome, and introns become the predominant site for the localization of such stretches because of their extensive length (21-fold longer than exons, Piovesan et al. 2016). These polyA stretches present additional priming sites (besides the 3’ polyA tail) for the polyT RT primer, thereby enabling more efficient transcript capture. Conversely, most intronic reads are therefore derived from genes with multiple polyA stretches (long genes), thereby introducing technical detection bias. This bias gets further magnified in snRNA-seq libraries that are inherently enriched for nascent transcripts (and hence intronic reads), and filtering such reads would mean reduced gene detection sensitivity, shallower sequencing depth and under-utilized sequencing cost. Here, we provided a normalization strategy for UMI counts derived from intronic reads that can remove gene-length associated technical biases. Implementation of this normalization strategy removes technical artifacts while retaining true biological features, thereby improving integration and enabling joint analysis of scRNA-seq and snRNA-seq datasets. In such joint analysis, our normalization strategy would also improve the accuracy of differential expression testing between any technique-specific clusters identified.

Finally, we demonstrated applicability of scVI for integration of scRNA-seq and snRNA-seq datasets. This is critical for the generation of a comprehensive adipose tissue atlas, since investigations into the stromal vascular fraction heterogeneity have been performed using scRNA-seq whereas snRNA-seq is favorable for investigations into the existence of adipocyte subtypes. Therefore, any efforts to identify shared subpopulations across such datasets, and the lineages therein would demand data integration. Overall, snRNA-seq provides an effective method for characterizing cellular heterogeneity and functionally relevant gene expression profiles within human preadipocytes and adipocytes. We expect that snRNA-seq will be actively adopted by the adipose community for high-throughput transcriptomic profiling of the tissue and aid in increasing its representation in initiatives such as the Human Cell Atlas. Ultimately, joint analysis of datasets acquired using multiple sequencing techniques will aid in the creation of a comprehensive human adipose tissue atlas, thereby enabling us to dissect its critical role in health and disease.

## 4 Methods

### 4.1 Preadipocyte culture and adipogenic differentiation

Detailed protocol for maintenance, cryopreservation, and differentiation of white and brown preadipocytes are outlined in a different study (Shamsi and Tseng 2017). Briefly, for culturing preadipocytes, cells were grown in DMEM medium (Corning, 10-017-CV) supplemented with 10% vol/vol FBS and containing 1% vol/vol Penicillin-Streptomycin (Gibco). Cell culture was maintained at 37°C in a humidified incubator containing 5% vol/vol CO2. 80% confluent cells were passaged using 0.25% trypsin with 0.1% EDTA (Gibco, 25200-056) for a 1:3 split in a new 100 mm cell culture dish (Corning).

Prior to adipogenic differentiation, white preadipocytes were allowed to grow up to 100% confluence in a 100 mm cell culture dish (Corning). After 48 hours at 100% confluence, growth media was replaced with adipogenic induction media every 48 hours for the next 20 days. Induction media was prepared by adding 1 mL FBS, 500 μl Penicillin-Streptomycin, 15 μl human Insulin (0.5 μM, Sigma-Aldrich, I2643-50MG), 10 μl T3 (2 nM, Sigma-Aldrich,T6397-100MG), 50 μl Biotin (33 μM, Sigma-Aldrich, B4639-100MG), 100 μl Pantothenate (17 μM, Sigma-Aldrich, P5155-100G), 1 μl Dexamethasone (0.1 μM, Sigma-Aldrich, D2915-100MG), 500 μl IBMX (500 μM, Sigma-Aldrich, I7018-100mg), and 12.5 μl Indomethacin (30 μM, Sigma-Aldrich, I7378-5G) to 48.5 mL DMEM medium and sterile filter.

### 4.2 Harvesting preadipocyte and mature adipocyte for scRNA-seq

At preadipocyte stage, cells were harvested from 100 mm plates, labeled with hashtag antibodies (Table S1A, Note S1), and finally suspended in PBS with 0.04% BSA at ∼1000 cells/uL concentration for downstream sequencing. At mature adipocyte stage, cells were first washed with PBS (Corning, 21-040-CV) and incubated with a monolayer of .25% trypsin with 0.1% EDTA (Gibco; 25200-056; monolayer obtained by adding and removing 1 mL of trypsin) for 2-3 minutes in a tissue culture incubator. When adipocytes started to become round and detached from the plate, trypsin was neutralized by adding 1 mL of FBS. Clumps of adipocytes were dislodged using a wide bore 1 mL pipette tip and filtered using a 70 μm cell strainer. Concentration of adipocyte suspension was adjusted to ∼ 200 cells/uL using FBS for downstream spotting using the CellenOne X1 machine.

### 4.3 Nuclei isolation from preadipocytes and mature adipocytes for snRNA-seq

Nuclei were isolated from white and brown preadipocytes using an NP-40 based lysis buffer: To 14.7 mL nuclease-free water (Qiagen), 150 ul of Tris-Hydrochloride (Sigma, T2194), 30 uL of Sodium Chloride (5M; Sigma, 59222C), 45 uL of Magnesium Chloride (1M; Sigma, M1028), and 75 uL of NP-40 (Sigma, 74385) was added. Two 100 mm dishes were used for nuclei isolation from each preadipocyte type. 500 uL of NP-40 based lysis buffer was added to each 100 mm dish and a cell scraper was employed to release adherent cells from the plates. Cells were then incubated with the lysis buffer for 5 minutes on ice in a pre-chilled 15 mL falcon tube. Cells were washed with ice-cold PBS supplemented with .2 U/uL RNase Inhibitor (Protector RNase Inhibitor; henceforth called wash buffer) 4 times by centrifuging at 500 rcf for 5 minutes at 4°C. Wash buffer was aspirated after the final round of centrifugation and nuclei were resuspended in the ice-cold wash buffer and filtered using a 40 um cell strainer. Final concentration was adjusted to ∼ 1000 nuclei/uL using a hemocytometer for downstream sequencing. Nuclei were also stained using 0.08% trypan blue dye to assess nuclear membrane integrity under brightfield imaging. For nuclear isolation at the mature adipocyte stage, the same protocol was implemented as mentioned above with the modification of using 1 mL lysis buffer for each 100 mm dish.

### 4.4 Single-cell and single-nuclei sequencing

**Table 1:** Sequencing metrics for individual libraries used in our study.

| Technique | Sample | Chemistry | Depth (per cell) | # cells | Sequencer |
| --- | --- | --- | --- | --- | --- |
| scRNA-seq | White & brown preadipocytes | 10X Chromium 3' v2 + hashtag antibodies | 78,000 | 8,000 | NovaSeq S2 |
| scRNA-seq | White adipocytes | mcSCRB-seq | 100,000 | 200 | NextSeq |
| snRNA-seq | White preadipocytes | 10X Chromium 3' v3 | 174,738 | 8,000 | NovaSeq S2 |
| snRNA-seq | Brown preadipocytes | 10X Chromium 3' v3 | 216,700 | 7,000 | NovaSeq S2 |
| snRNA-seq | White adipocytes | 10X Chromium 3' v3 | 123,700 | 12,000 | NovaSeq S2 |

For mcSCRB-seq experiment with white adipocytes (day 20), 96-well plates were first preloaded with rows of 10 uniquely barcoded primers and lysis buffer according to the mcSCRB-seq protocol, with the only difference being the use of μCB-seq RT primers (Chen et al. 2020) instead of standard mcSCRB-seq ones. The sequence of barcodes used were: TCACAGCA, GTAGCACT, ATAGCGTC, CTAGCTGA, CTACGACA, GTACGCAT, ACATGCGT, GCATGTAC, ATACGTGC, and GCAGTATC. CellenONE X1 instrument was used to individually deliver a single adipocyte into each well for a total of 200 cells. Following cell delivery, the mcSCRB-seq protocol was followed directly, but with the following two modifications:

1. A 1:1 ratio of AmPure XP beads was used to pool all cDNA after RT as opposed to the manual bead formulation from standard mcSCRB-seq
2. NEBNext i5 indexed primers (NEB, E7600 and E7645) were used as opposed to the non-indexed P5NEXTPT5 primer during library PCR and indexing step to generate dual indexed libraries for multiplexing

### 4.5 scRNA-seq and snRNA-seq data analysis

scRNA-seq white & brown preadipocytes dataset was processed using cellranger-3.0.2 with default parameters, and the human GRCh38-3.0.0 genome (November 19, 2018) as input. A custom pre-mRNA GTF file was created using the GRCh38-3.0.0 FASTA file as input to include intronic reads in UMI counts. Sample demultiplexing, doublet removal, and empty droplet removal was performed using the Seurat (Butler et al., 2018) function *HTODemux* (Note S1). Cell barcodes were further filtered to have more than 200 genes. Post demultiplexing and filtering, scVI (Lopez et al. 2018) was used to infer a 20-dimensional latent space based on the expression of the top 2000 most variable genes. This latent space was then used in Seurat to generate the UMAP visualization using the RunUMAP command. Downstream clustering (resolution = 0.4) and differential expression analysis (logFC > 0.5) was performed using Seurat’s SCTransform pipeline (Hafemeister and Satija 2019). Clusters with > 5% mean mitochondrial content were removed from downstream analyses. Gene ontology analysis was performed at geneontology.org (Ashburner et al. 2000; Carbon et al. 2019; Mi et al. 2019) and results were further confirmed using the goana package in R with genome wide human annotation derived from org.Hs.eg.db Bioconductor package. Transcription factor enrichment analysis was performed using the ChEA3 tool (Keenan et al. 2019). GRCh38-ref20202A (2020) reference was used for analysis involving lncRNAs, keeping everything else the same.

snRNA-seq white and brown preadipocyte dataset was also processed using cellranger-3.0.2. For white preadipocyte, barcodes with < 200 genes were removed from downstream analyses. CellBender (Fleming et al. 2019) was used to remove empty droplets. For downstream analyses, only barcodes called as cells by both cellranger and CellBender were used and barcodes with UMI count > 49000 were filtered out as possible doublets. For brown preadipocyte, barcodes with < 200 genes were removed and scVI was used to infer a 20-dimensional latent space. First round of clustering was performed in Seurat with the resolution set to 0.06. We identified 3 clusters, with cluster 1 having most of the barcodes called as empty by CellBender. Therefore, cluster 1 was removed from downstream analysis as well other barcodes that were called as “cell-containing” by cellranger but not by CellBender. Cluster 2 was marked with high mitochondrial content (>20%) and hence was also removed from downstream analyses. After filtering out low-quality barcodes and clusters, Scrublet (Wolock et al. 2019) was used to remove any potential doublets. After individual QC of white and brown preadipocyte libraries, the two datasets were integrated together using scVI with no batch effect correction. The output from scVI analysis was a 20-dimensional latent space representation with cell embeddings for both white and brown nuclei. This latent space was then used in Seurat to generate the UMAP visualization using the RunUMAP command. Downstream clustering (resolution = 0.24) and differential expression analysis (logFC > 0.5) was performed using Seurat’s SCTransform pipeline (see Fig. S4). For gene ontology, and differential expression analyses, the same tools as mentioned in the above paragraph were used. GRCh38-ref20202A (2020) reference was used for analysis involving lncRNAs, keeping everything else the same.

mcSCRB-seq white adipocyte dataset was processed using zUMIs (Parekh et al. 2018) using the GRCh38 index for STAR alignment. We provided the 10X CellRanger recommended GRCh38-3.0.0 GTF file as input for standardization of gene counts. Reads with any barcode or UMI bases under the quality threshold of 20 were filtered out and known barcode sequences were supplied in an external text file. UMIs within 1 hamming distance were collapsed to ensure that molecules were not double-counted due to PCR or sequencing errors. Only exonic reads were counted towards UMI quantification. The umi-count matrix generated using zUMIs was read using the readRDS command in Seurat. The CellenOne X1 machine acquires an image of every cell spotted and the presence of a single cell was further validated by analyzing these images to remove possibly empty or doublet barcodes. The Seurat object was analyzed using a standard Seurat pipeline with resolution set to 0.6 for clustering.

snRNA-seq white adipocyte dataset was processed using cellranger-3.1.0. Barcodes with < 200 genes were removed from downstream analyses and scVI was used to infer a 20-dimensional latent space. For clustering using Seurat, the resolution parameter was set to 0.45. We identified 7 clusters, with cluster 3 having most of the barcodes called as empty by CellBender. Therefore, cluster 3 was removed from downstream analysis as well other barcodes that were called as “cell-containing” by cellranger but not by CellBender. Cluster 5 was marked with high mitochondrial content and hence was also removed from downstream analyses. Cluster 2 had the greatest number of doublets identified by the doubletDetection (Gayoso et al. 2019) tools and was filtered out, as well as cluster 4 which was enriched for ribosomal proteins suggesting cellular debris contamination.

### 4.6 Identifying number of lncRNAs detected as a function of sequencing depth

For identifying the number of lncRNAs detected as a function of sequencing depth, the fastq files for scRNA-seq preadipocyte dataset only were subsampled using seqtk v1.3 with the random seed = 100. For each subsample depth, fastq files were processed using cellranger-3.1.0 with GRCh38-ref2020A pre-mRNA as the reference. snRNA-seq data for white and brown nuclei (as processed with cellranger at full depth) were then aggregated with the output of scRNA-seq preadipocyte data at varying sequencing depth using the cellranger aggr command to achieve same number of average transcriptome mapped reads. Number of lncRNAs detected were then calculated as a function of sequencing depth, with lncRNA assumed as detected in a given cell/nuclei if UMI count >0.

### 4.7 Silhouette coefficient analysis

Both scRNA-seq and snRNA-seq datasets for brown preadipocytes were subsampled as described above. snRNA-seq dataset was further randomly subset to have the same number of total barcodes as scRNA-seq. At each sequencing depth, top 20 principal components were calculated using Seurat’s standard pipeline. Three resolution coefficients based on the Silhouette index, Calinski Harabasz index, and Davies Bouldin index were then calculated based on Euclidean distance between cells in the PCA space using the clusterCrit package in R. For analyzing cluster separation resolution between brown cluster 1 and 2 as a function of UMI count, exactly the same analysis was performed except that downsampling was performed to have the same number of UMI rather than reads between scRNA-seq and snRNA-seq dataset using the downsampleMatrix command in the DropletUtils package in R (Lun et al. 2019).

### 4.8 Integration of snRNA-seq and scRNA-seq data

For integrating scRNA-seq white preadipocyte (day-0) & white-adipocyte (day-20) and snRNA-seq white preadipocyte (day-0) & white-adipocyte (day-20) datasets (a total of 4 datasets), we first created a single anndata object with UMI count-matrices from each dataset as input. Each of the four UMI matrices were generated by processing the originals fastq files (no downsampling of reads), and subset to only have high-quality barcodes as outlined in Methods section # 5. During concatenation, each of the four datasets was assigned a “batch” key. The concatenated anndata object was then used as input to scvi-tools for integration using the commands outlined in the tutorial here: https://docs.scvi-tools.org/en/stable/user_guide/notebooks/harmonization.html. The output of following these steps was a 10-dimensional latent space with batch-corrected embedding for cells from each of the four datasets. UMAP visualization was then generated using the RunUMAP command in Seurat with the 10-dimensional latent space as input. The dendrogram was generated using the *BuildClusterTree command* in Seurat, which constructs a phylogenetic tree relating the ‘average’ cell from each identity class. Tree is estimated based on the eigenvalue-weighted euclidean distance matrix constructed in latent-dimension space.

## 5 Data Access

Data related to this study is available upon request to the corresponding author.

## 7 Acknowledgments

This publication was supported by the National Institute of General Medical Sciences of the National Institutes of Health under award number R35GM124916. This publication was also supported by Grant No. CZF2019-002454 from the Chan Zuckerberg Foundation. Aaron Streets is a Chan-Zuckerberg Biohub Investigator and a Pew Scholar in the Biomedical Sciences, supported by the Pew Charitable Trusts. AG is supported by the UC Berkeley Lloyd Fellowship in Bioengineering.

