## Supplemental Data for "Characterization of transcript enrichment and detection bias in single-nuclei RNA-seq for mapping of distinct human adipocyte lineages"

**Note S1: Hashing of white and brown preadipocytes using oligo-conjugated hashtag antibodies**

Cell hashing enables pooling of all samples prior to loading them onto a single 10X chromium controller lane, thereby enabling combined library preparation and sequencing of all samples to eliminate potential "batch" artifacts. During single-cell suspension preparation of white and brown preadipocytes for downstream scRNA-seq, we split each preadipocyte type into two individual microcentrifuge tubes for a total of four working samples. Brown preadipocytes were labelled with Hashtag-A0251 and A0252 antibodies, and white preadipocytes with A0253 and A0254 antibodies (Table S1A). By sequencing these hashtag antibodies alongside the cellular transcriptome, we assigned each cell to its sample of origin, and identified doublets originating from multiple samples. Hashtag-antibody library was counted using the CITE-seq-Count workflow (10.5281/zenodo.2585469) and demultiplexed using the Seurat function 'MULTIseqDemux'. Demultiplexing pipeline identified 143 negative barcodes, 532 doublet barcodes, and 6918 cell-containing barcodes. As expected, every cell-containing barcode had highly positive and specific expression of only a single hashtag antibody, every doublet had marked expression for a combination of two antibodies, and negatives had very low expression for all antibodies (Fig. S2A).

**Labeling protocol:** For staining cells with hashtag antibodies, we followed supplier's protocol (https://www.protocols.io/view/cell-hashing-nfzdbp6). Briefly, cells were harvested from a single 100 mm cell-culture dish and suspended in 100 µl of cell staining buffer in 2 ml low bind tubes. 5 µl of Human TruStain FcX™ Fc Blocking reagent was then added, and cells were incubated for 10 minutes at 4°C. 0.5 ug of a unique Cell Hashing antibody was added to each tube and cells were incubate for 30 minutes at 4°C. Cells were washed with 1 mL of cell staining buffer for 3 times by centrifuging at 1200 rpm for 4 minutes at 4°C. Finally, cells were suspended in PBS and 0.04% BSA at ~ 1000 cells/uL for downstream 10X sequencing.

**Note S2: Proliferating vs growth arrested cells in snRNA-seq and scRNA-seq white preadipocyte dataset**

As highlighted in Fig. 6B UMAP visualization, both scRNA-seq and snRNA-seq white preadipocyte dataset (day-0) were cleaved into two halves. We investigated the differences between these two halves by manually annotating clusters as following (Fig. S8A):

Cluster 0: day-20-differentiating-preadipocyte nuclei and cells

Cluster 1: day-20-adipocyte nuclei and cells

Cluster 2: top half of cleaved day-0-preadipocyte nuclei and cells

Cluster 3: bottom half of cleaved day-0-preadipocyte nuclei and cells

Cluster 4: day-20-cluster-2-nuclei

We normalized the data using *NormalizeData* command in Seurat and plotted expression profiles of proliferation and mitotic marker genes *PLK1*, *MYBL2*, *BUB1*, *MKI67*, *CDK1*, and *CCNB1* (Fig. S8B to S8G). As expected, cluster 0 and 1 which primarily comprised of day-20 cells had no expression of proliferation markers, which is in line with their growth arrested behavior post adipogenic induction. However, prior to differentiation (day-0-preadipocytes), cells undergo cell cycle progression, thereby explaining the positive expression of proliferation marker genes in cluster 3. Cluster 2 cells were perhaps preadipocytes that underwent growth arrest due to contact inhibition during cell culture. Notably, even after 20 days of differentiation, a very small number of cells (cluster 4) were still highly proliferating, suggesting that these cells could be preadipocytes that never underwent growth arrest. We also calculated cell cycle phase scores based on canonical markers using the Cell-Cycle Scoring pipeline in Seurat and assigned either G1, G2M, or S Phase to each of these cells. As expected, most of the cells in clusters 0, 1, and 2 were in G1 phase as opposed to G2M and S phase in clusters 3 and 4 (Fig. S8H).

Note S3: Outline of normalization strategy to correct for gene-length-based detection bias arising from including intronic reads

1. Rationale for Normalization

Multiple recent studies have demonstrated internal hybridization of polyT RT-primer to intronic polyA stretches as the primary mechanism for the capture and detection of intronic reads (La Manno et al. 2018; Patrick et al. 2020; Shulman and Elkon 2019). Assuming that all intronic reads are derived from such hybridization incidences, number of observed **intronic** UMIs for any gene $g$ in a given nuclei can be estimated by the following equation:

$$p_i \times pA_g \times N_g = x_g \qquad \textit{Equation 1}$$

where $p_i$ is the probability to capture an intronic read, $pA_g$ are the number of polyA stretches in gene $g$ and $N_g$ is the true transcript abundance. Assuming that $p_i$ is independent of gene $g$

$$N_g \propto \frac{x_g}{pA_g} \qquad \textit{Equation 2}$$

2. Estimating $\boldsymbol{pA_g}$

$pA_g$ can be modeled by the following equation, where $pd$ is the number of polyA stretches per kilobase of the genic region in the human genome, and $gl_g$ is the total length of the gene in kilobase, including introns and exons.

$$pA_g = pd \times gl_g \qquad \textit{Equation 3}$$

Since the polyT tail in 10x Chromium RT primer is 30-bp long, we assumed hybridization to occur between the polyT tail and a polyA stretch, if the polyA sequence is at least 15-bp long (50% of the polyT tail). We queried the GRCh38 human genome to get positions of all polyA tracts at least 15-bp long, without mismatch, and screened for overlaps between such polyA tracts and gene coordinates for all genes in the cellranger GRCh38-2020A reference (which includes lncRNAs). As expected, total number of polyA tracts were highly correlated with gene length (Spearman R = 0.82, p-value < 0.05, Fig.

S1A) for each gene. We also calculated mean number of polyA tracts per Kbp for each gene, and estimated $pd$ as the mean number of polyA tracts per Kbp across all genes, including zeroes (Fig. S1B). Following this analysis, we estimated $pd$ to be equal to 0.07.

We retrieved gene coordinates, strand, and gene length information using the GRCh38 gene annotation file downloaded from Gencode (Release 32). The same GTF was used for cellranger analysis. Briefly, each gene was first summarized by setting 3rd column in the GTF to **gene**, followed by calculation of gene length by subtracting the 5th and 4th columns.

### 3. Normalization Strategy

Based on *Equation 2*, we present a normalization framework to reduce the technical bias arising from comparisons of nuclear and cellular data upon inclusion of intronic reads. This normalization strategy is implemented on the count matrix generated using **only intronic reads**, for both scRNA-seq and snRNA-seq datasets, and provides a modified UMI count-abundance, taking gene-length into account, for each cell and nuclei, based on the following equations:

$$\overline{x_g} = \frac{x_g}{gl_g \times pd} \qquad \textit{Equation 4}$$

where $x_g$ is the original UMI-count for gene $g$ in a given cell/nuclei, and $\overline{x_g}$ is the modified UMI count after normalization for gene $g$ in the same cell/nuclei. This modified intronic UMI-count is then added to the observed exonic UMI-count for each gene $g$ in a given cell/nuclei and finally library-normalized as following:

$$\bar{z_g} = \log\left(\frac{\overline{x_g}+y_g}{N_i+N_e} \times e^4 + 1\right) \qquad \textit{Equation 5}$$

$$N_i = \sum_{g \in G} \overline{x_g} \qquad \textit{Equation 6}$$

$$N_e = \sum_{g \in G} y_g \qquad \textit{Equation 7}$$

where $y_g$ is the original UMI-count for gene $g$ using exonic reads, and $\bar{z_g}$ is the final log-normalized count used for downstream differential expression testing between cells and nuclei.

**Note S4: Classification of differentially expressed genes in scRNA-seq dataset as marker genes**

Fluorescent-activated cell sorting (FACS) has been instrumental in identifying lineage-specific preadipocyte marker genes in mice (Hepler et al. 2017). However, markers identified in mice are not comprehensively selective for humans (de Jong et al. 2015; Ferrero et al. 2020). We therefore sought to define a set of white-specific and brown-specific marker genes as well as a set of genes specifically expressed in cluster 1 and cluster 2. Using the identified list of differentially expressed genes (Table S1B and S1C), we implemented stringent cutoff criteria with logFC > .8 in each cell-type, minimum detection of 60 % and maximum detection of 40% in the other cell-type, for classifying genes with highly enriched and specific expression as marker genes. On this basis, we recognized *NTNG1*, *RPL39L*, *PGF*, *LAMA4*, *BAALC*, *HIP1*, and *HAS2* as markers of white preadipocytes; *LIMCH1*, *LYPD1*, *RGS4*, *ITGBL1*, *CDH13*, and *COL4A2* as markers of brown preadipocytes in the human neck depot (Fig. 1B highlighted in beige and Table S1B). We also identified *KRT18*, *LUZP2*, *DLGAP1*, *SBSPON*, *MAP3K7CL*, and *NRXN3* as markers of brown cluster 1; *CTSK*, *BST2*, and *MOXD1* as markers of brown cluster 2 (Fig. 1C highlighted in beige and Table S1C).

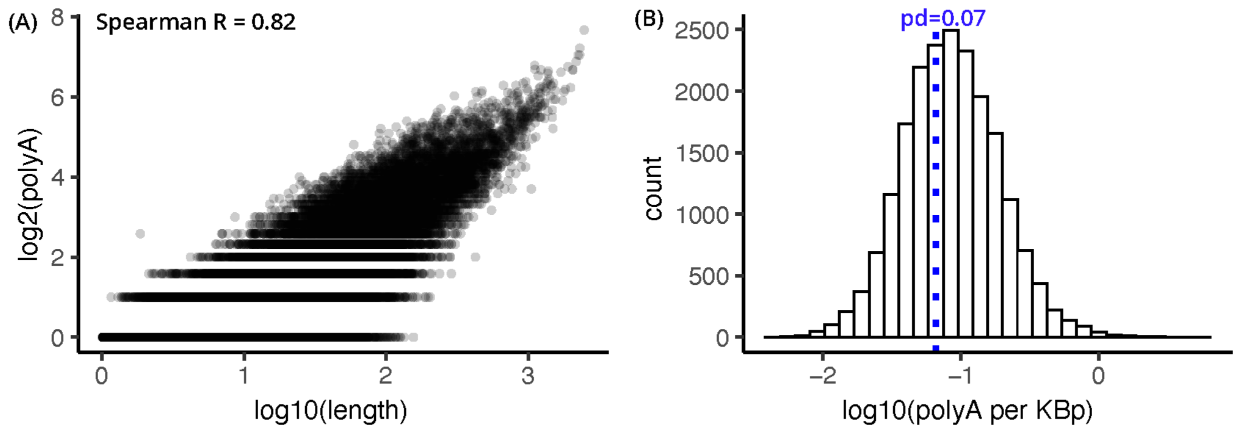


**Figure S1 (related to Note S3) Estimating polyA-tract density per Kbp in the genic region. (A)** Scatter plot of total number of polyA-tracts (greater than 15-bp) plotted against gene length. Each dot is a gene in the GRCh38-2020A reference from cellranger analysis pipeline. **(B)** Distribution of mean number of polyA-tracts per Kbp for each gene in panel A. Blue dotted line indicates mean number of poly-A tracts per Kbp across all genes and is used to estimate *pd*=0.07. See NoteS3 for details on normalization strategy.

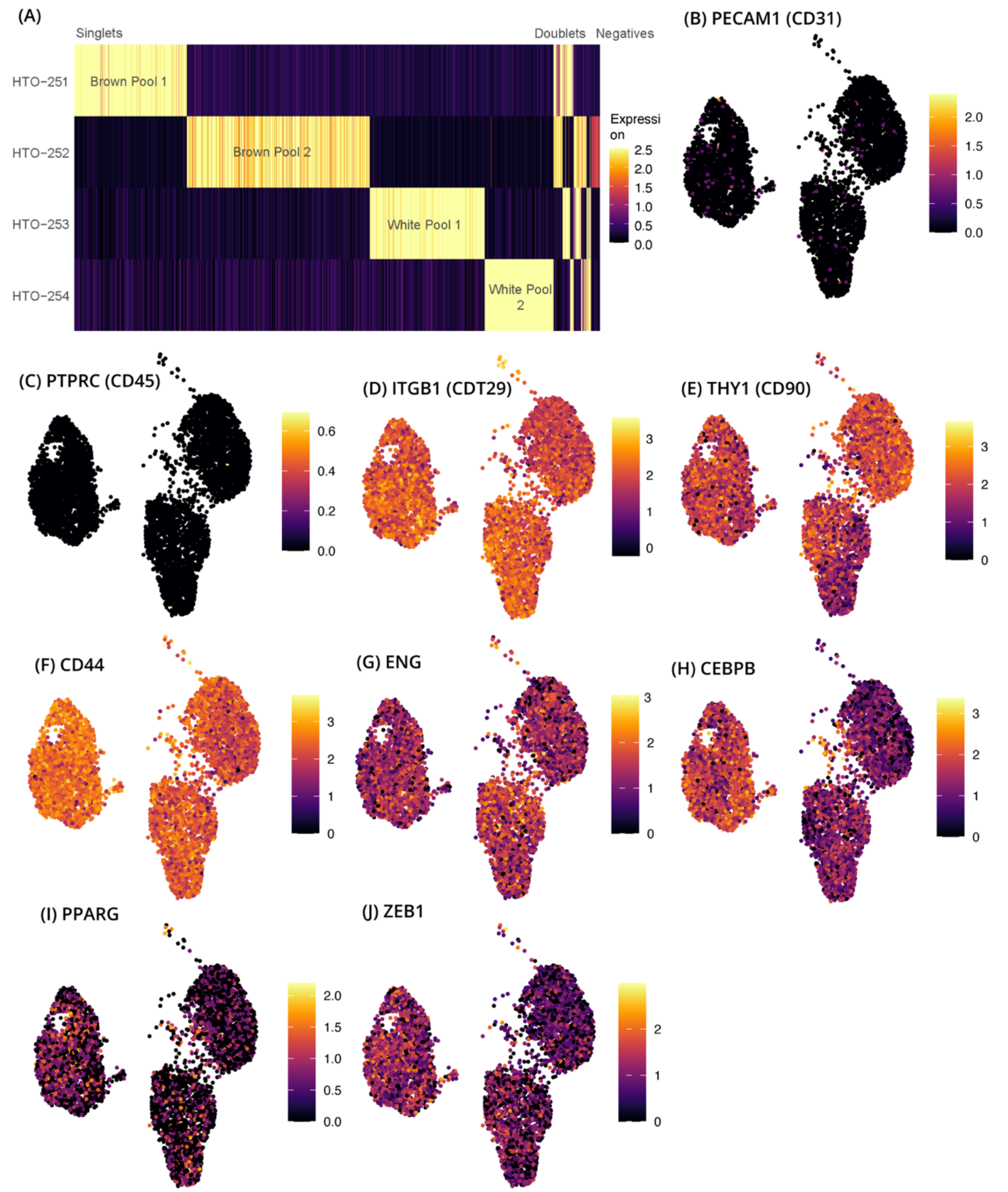


**Figure S2 (related to Result # 1 and Figure 1) Analysis of white and brown preadipocyte scRNA-seq dataset (A)** Log-normalized expression of four hashtag antibodies used for multiplexing of white and brown preadipocytes (whole-cells). Each row marks the expression of a given antibody in 5000 randomly sampled barcodes (columns). Also see Note S1 and Table S1A. **(B) to (J)** Expression profiles of marker genes in scRNA-seq dataset.

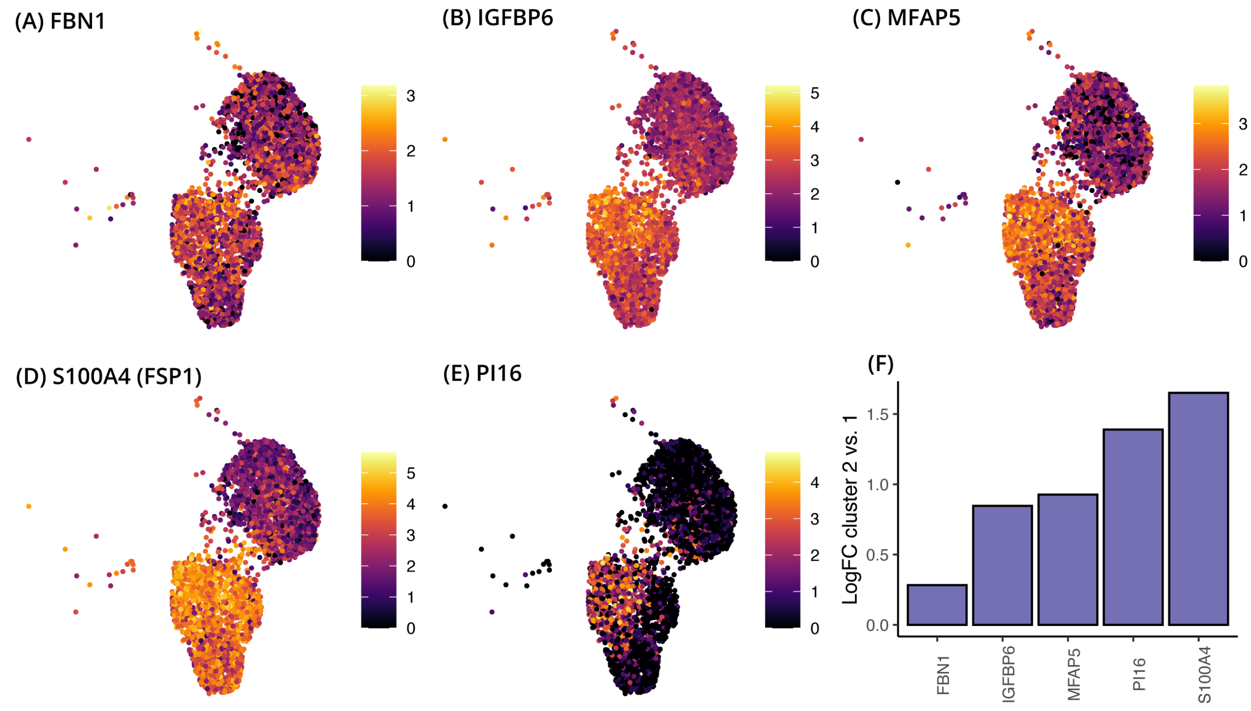


**Figure S3 (related to Result # 1 and Figure 1) Differential expression in brown preadipocyte scRNA-seq dataset between cluster 2 and cluster 1. (A) to (E)** Expression profile of marker genes for Fsp1+ fibroblasts identified in (Vijay et al. 2020) **(F)** Log fold change values of the marker genes as calculated using cluster 2 vs cluster 1 differential expression test. All genes were significantly enriched in cluster 2 with FDR < 0.05. Also see Table S1C.

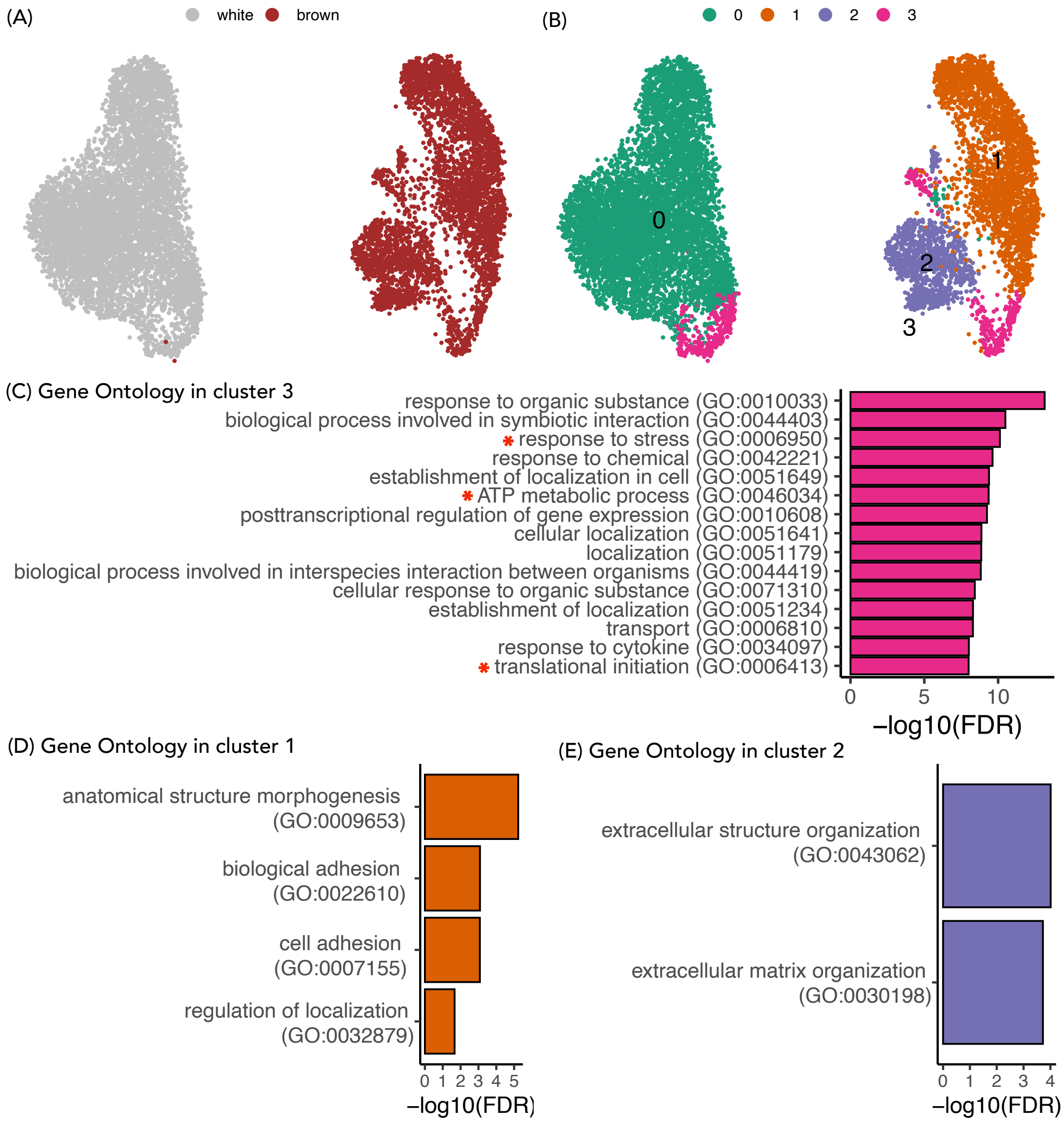


**Figure S4 (related to Result # 2 and Figure 2) Unsupervised clustering of white and brown preadipocytes snRNA-seq dataset (A) and (B)** UMAP visualization of white and brown preadipocytes annotated either manually to reflect the sample of origin (A) or based on unsupervised clustering (B). **(C)** Top gene ontology biological processes (BP) terms enriched in cluster 3 based on a cluster 3 vs. all DE test. Marked in red is the enrichment of BP terms because of stress response genes (response to stress), mitochondrial genes (ATP metabolic process), and ribosomal mRNA genes (translational initiation). Enrichment of mitochondrial and ribosomal mRNA genes indicates the presence of cellular background RNA contamination (see Fig. S5). **(D)** Top 10 gene ontology terms in brown cluster 1 in scRNA-seq dataset (Fig. 1D) that are also enriched in cluster 1 in snRNA-seq dataset. None of these top 10 GO terms were enriched in cluster 2 in snRNA-seq dataset. **(E)** Top 10 gene ontology terms in brown cluster 2 in scRNA-seq dataset (Fig. 1D) that are also enriched in cluster 2 in snRNA-seq dataset. None of these top 10 GO terms were enriched in cluster 1 in snRNA-seq dataset.

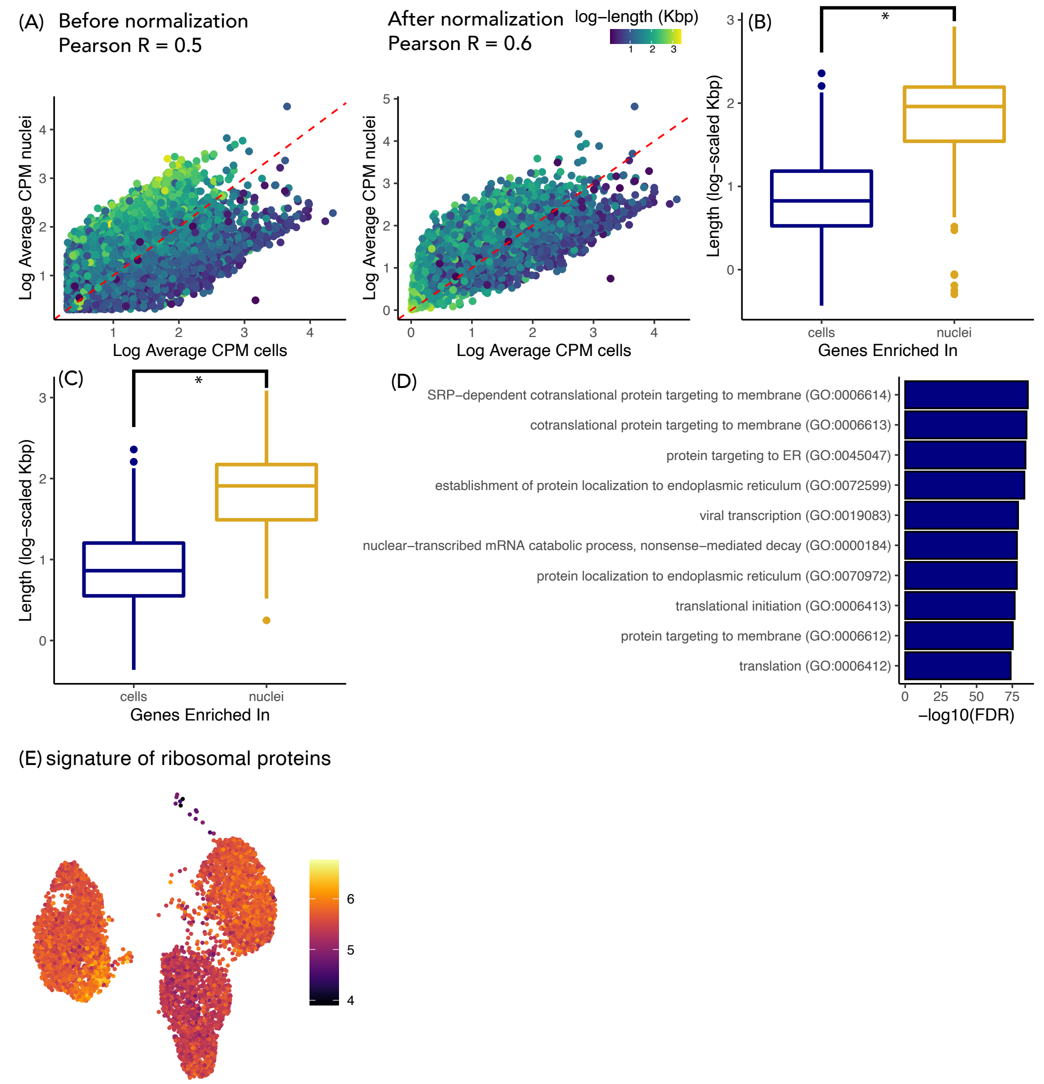


**Figure S5 (related to Result # 3 and Figure 3) Gene length-associated detection bias in snRNA-seq. (A)** Average expression of genes in white cells and white nuclei when using both intronic and exonic reads, without normalization (left panel), and with normalization (right panel). See Note S3 for normalization strategy. Each dot represents a gene with average counts per million (CPM) > 1 when using both intronic and exonic reads, without normalization, in both cells and nuclei. White nuclei were randomly selected to have as many barcodes as white cells. Red dotted line has slope = 1. **(B)** Distribution of gene length for genes enriched in cells (in blue) and nuclei (in yellow) with log fold-change > 1 and FDR < 0.05 including both intronic and exonic reads. Intronic UMI-count matrix was normalized to correct for gene length bias in both cells and nuclei (see Note S3). **(C)** Distribution of gene length for genes enriched in cells (in blue) and nuclei (in yellow) with log fold-change > 1 and FDR < 0.05 using only exonic reads. **(D)** Top 10 gene ontology terms enriched in white cells as compared to white nuclei based on differential expression after normalization. **(E)** Heatmap of transcriptional signature score defined using top 100 genes enriched in cells vs. nuclei in white preadipocytes based on log fold-change values after normalization. The scores are plotted on the 2D UMAP visualization of scRNA-seq preadipocyte data.

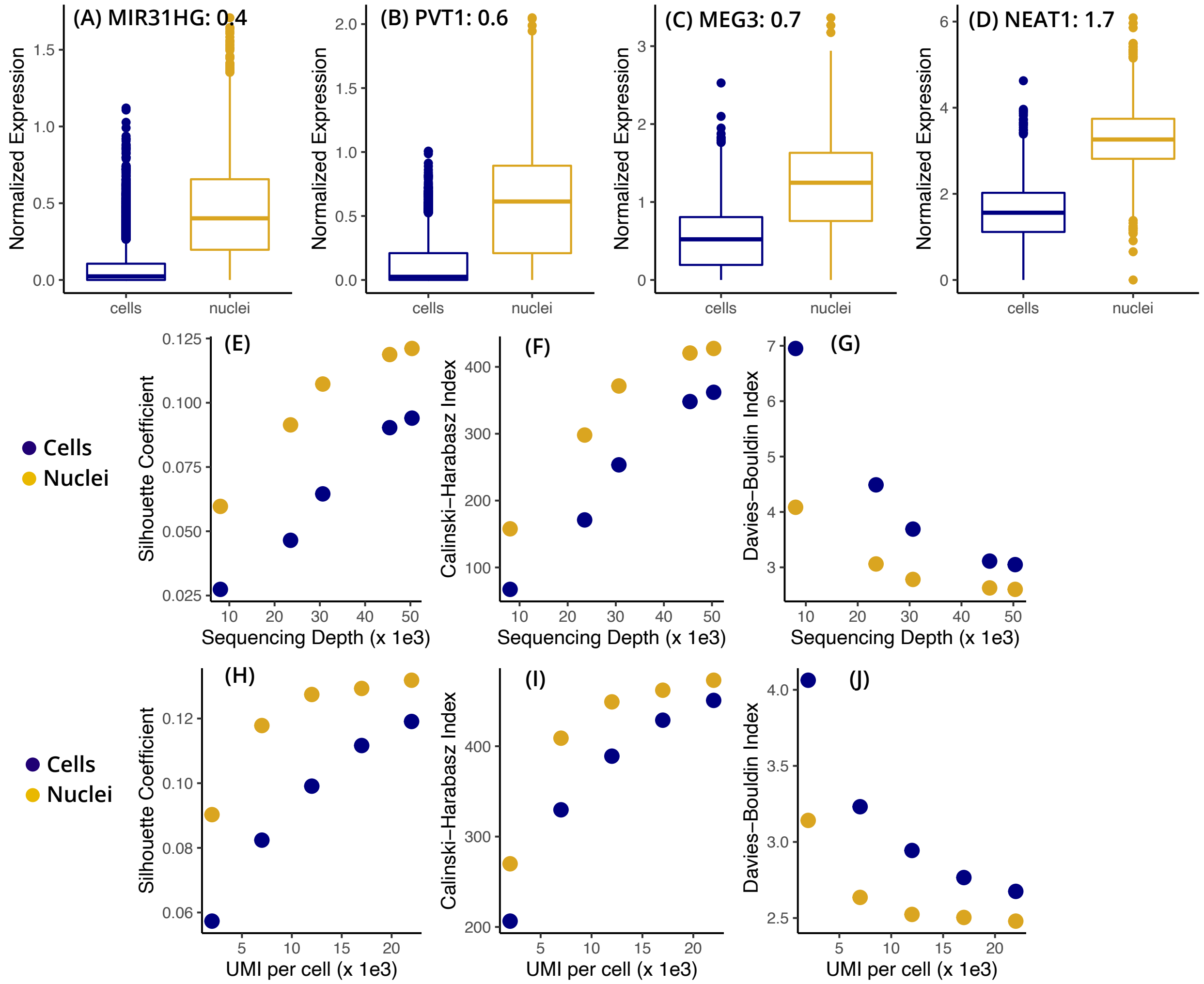


**Figure S6 (related to Result # 4 and Figure 4) Enrichment of lncRNAs in the nuclear transcriptome. (A) to (D)** Expression of adipogenic regulatory lncRNAs in brown nuclei over brown whole cells. Black text indicates logFC value for brown nuclei vs. brown cells DE test with FDR < 0.05 after normalization. **(E) to (G)** Cluster separation resolution quantification between brown cluster 2 vs cluster 1 in scRNA-seq and snRNA-seq dataset. **Only lncRNAs were considered for PCA manifold generation.** Both datasets were subsampled to have the same number of cells/nuclei and same number of mean transcriptome mapped reads. **(H) to (J)** Similar analysis as panel (E) to (G) but normalization was performed to have the same number of UMI counts per cell/nuclei. A higher Silhouette coefficient and Calinski Harabasz and a lower Davies Bouldin index indicate superior cluster separation performance.

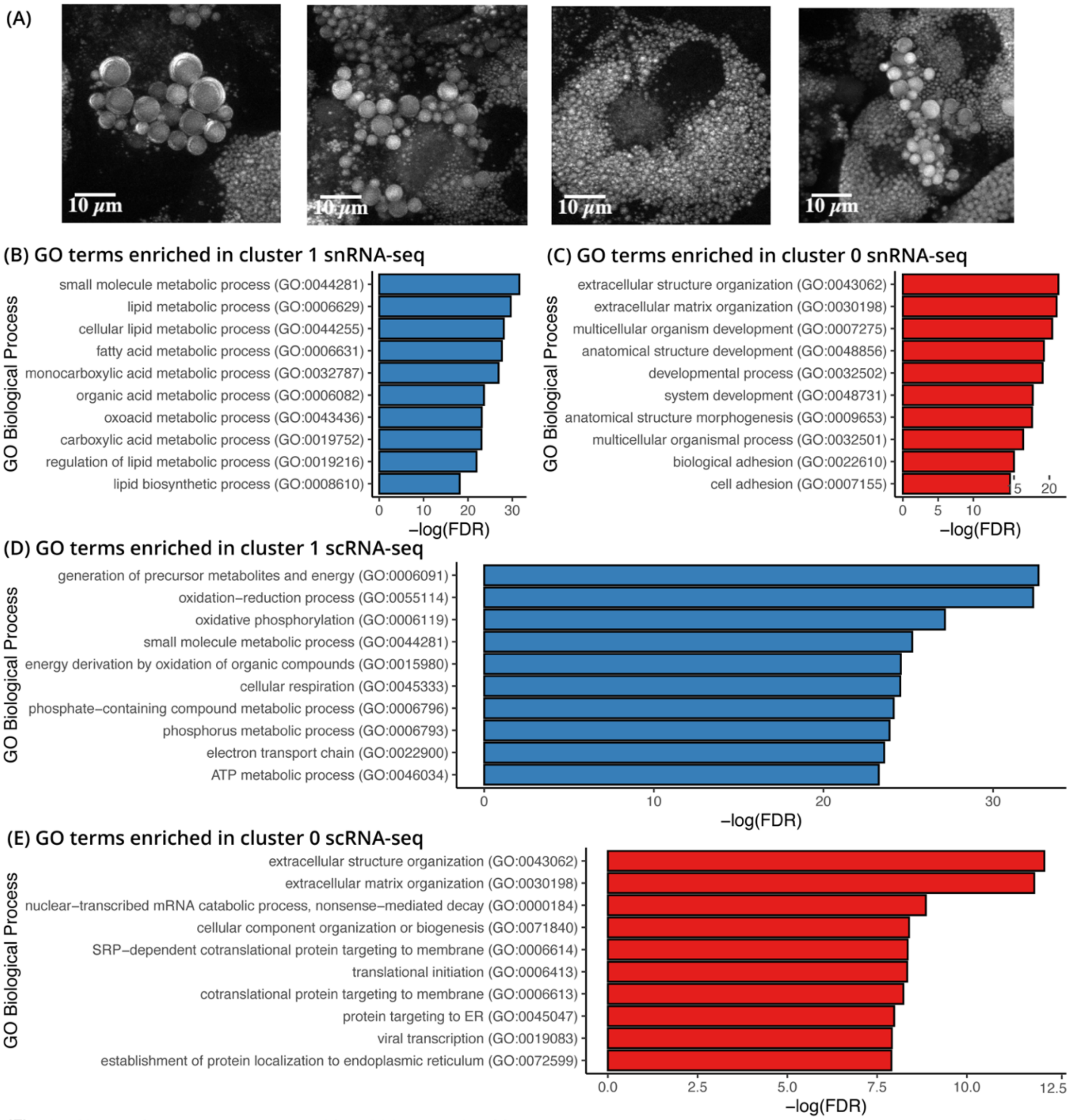


**(F)** DHCR24; HIST2H2AA4; UHMK1; MGST3; PRDX6; TIMP4; COX7A2;
NDUFA4; CD36; NDUFA1; UQCRB; COX6C; NDUFB9; STOM; COX8A; CRYAB; H2AFJ
COX14; ATP5F1B; COX6A1; AC126365.1; NDUFV2; ATP5F1A; ECH1; APOE; APOC1; UQCR10

**Figure S7 (related to Result # 5 and Figure 5) Comparative analysis of nuclear and whole-cell transcriptome at mature adipocyte stage (A)** Coherent anti-stokes Raman imaging of human white preadipocytes differentiated for 20 days using a chemical adipogenic induction cocktail. The images were acquired at 2845 $cm^{-1}$ wavenumber, which corresponds to the $CH_3$ peak present in lipids. Z-stacked images were acquired and the maximum intensity projection for each pixel was plotted. **(B) and (C)** Top 10 gene ontology terms enriched in cluster 1 (panel B) and cluster 0 (panel C) in **snRNA-seq dataset. (D) and (E)** Top 10 gene ontology terms enriched in cluster 1 (panel D) and cluster 0 (panel E) in **scRNA-seq dataset. (F)** List of 27 genes differentially enriched in cluster 1 (mature adipocytes) in scRNA-seq dataset but not differentially enriched in cluster 1 of snRNA-seq dataset

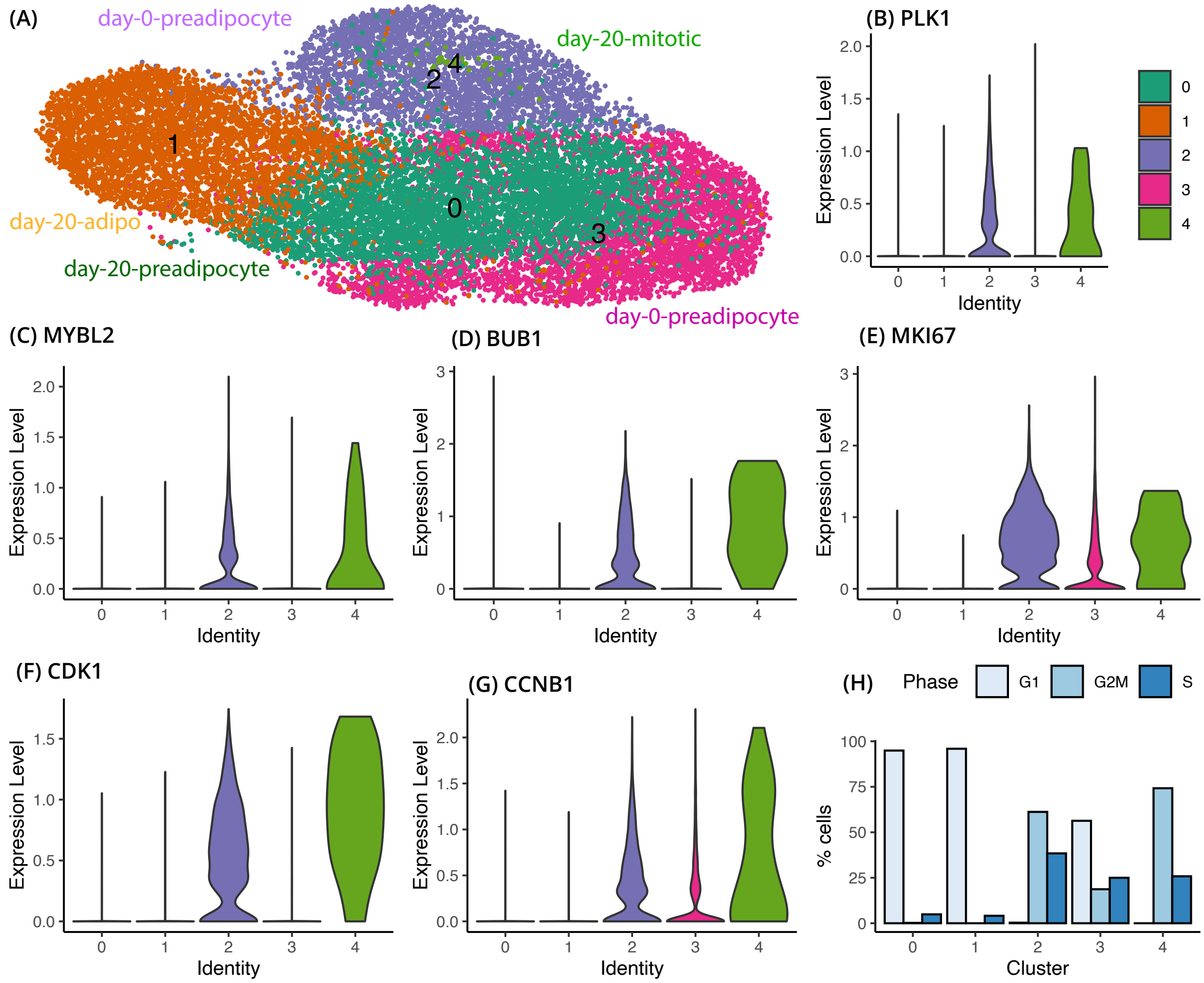


**Figure S8 (related to Note S2, Result # 6 and Figure 6): Proliferating vs growth arrested cells in snRNA-seq and scRNA-seq white preadipocyte dataset. (A)** Supervised clustering of integrated scRNA-seq and snRNA-seq white preadipocyte (day-0) and white adipocyte (day-20) dataset. See Note S2 for details regarding clustering scheme. **(B) to (G)** Violin plots of common proliferation and mitosis marker genes in clusters identified in panel (A). **(H)** Bar plot of distribution of cell cycle phase assignment in the clusters identified in panel (A). Y-axis plots the percent of cells belonging to different cell cycle phase for every cluster. See Note S2 for details regarding cell cycle phase assignment.
